# Oxidative Stress is a shared characteristic of ME/CFS and Long COVID

**DOI:** 10.1101/2024.05.04.592477

**Authors:** Vishnu Shankar, Julie Wilhelmy, Ellis J. Curtis, Basil Michael, Layla Cervantes, Vamsee A. Mallajosyula, Ronald W. Davis, Michael Snyder, Shady Younis, William H. Robinson, Sadasivan Shankar, Paul S. Mischel, Hector Bonilla, Mark M. Davis

## Abstract

More than 65 million individuals worldwide are estimated to have Long COVID (LC), a complex multisystemic condition, wherein patients of all ages report fatigue, post-exertional malaise, and other symptoms resembling myalgic encephalomyelitis / chronic fatigue syndrome (ME/CFS). With no current treatments or reliable diagnostic markers, there is an urgent need to define the molecular underpinnings of these conditions. By studying bioenergetic characteristics of peripheral blood lymphocytes in over 16 healthy controls, 15 ME/CFS, and 15 LC, we find both ME/CFS and LC donors exhibit signs of elevated oxidative stress, relative to healthy controls, especially in the memory subset. Using a combination of flow cytometry, bulk RNA-seq analysis, mass spectrometry, and systems chemistry analysis, we also observed aberrations in ROS clearance pathways including elevated glutathione levels, decreases in mitochondrial superoxide dismutase levels, and glutathione peroxidase 4 mediated lipid oxidative damage. Critically, these changes in redox pathways show striking sex-specific trends. While females diagnosed with ME/CFS exhibit higher total ROS and mitochondrial calcium levels, males with an ME/CFS diagnosis have normal ROS levels, but larger changes in lipid oxidative damage. Further analyses show that higher ROS levels correlates with hyperproliferation of T cells in females, consistent with the known role of elevated ROS levels in the initiation of proliferation. This hyperproliferation of T cells can be attenuated by metformin, suggesting this FDA-approved drug as a possible treatment, as also suggested by a recent clinical study of LC patients. Thus, we report that both ME/CFS and LC are mechanistically related and could be diagnosed with quantitative blood cell measurements. We also suggest that effective, patient tailored drugs might be discovered using standard lymphocyte stimulation assays.

## Introduction

With more than 660 million documented COVID cases worldwide, it has been estimated that as many as 65 million individuals may have “Long COVID”, a complex multisystemic condition associated with post-acute sequelae of Severe Acute Respiratory Syndrome Coronavirus 2 (SARS-CoV-2) infection (PASC)^1^. Long COVID spans all ages, affecting even those who have experienced only moderate COVID infections, with 10-12% vaccinated, 10-30% of non-hospitalized, 50-70% of hospitalized COVID-19 infection survivors estimated to endure persistent symptoms after infection^1,2^, with symptoms affecting individual quality of life and functional status^71^. Although the clinical presentation of Long COVID (LC) is highly heterogeneous, with adverse events spanning multiple organ systems from dysrhythmia and higher incidence of cardiac disorders^1,3^ to neurological and cognitive deficits such as memory loss and “brain fog”^2,4^, there are several common shared symptoms of LC, including fatigue (estimated pooled prevalence of 47%), shortness of breath (32%), and muscle pain (25%)^5,73^.

Strikingly, the clinical presentation of LC strongly resembles myalgic encephalomyelitis/chronic fatigue syndrome (ME/CFS), a complex chronic disease estimated to affect between 836,000 to 2.5 million individuals in the US alone^7^. By meta-analysis, it is estimated that 0.01 to 7.62% of the world population has ME/CFS, with 2x higher preponderance among females compared to males^68^. Based on three symptoms defined by the National Academy of Medicine criteria^7^, ME/CFS is characterized by (1) profound fatigue lasting for at least 6 months, (2) post-exertional malaise, and (3) unrefreshing sleep^6-8^. Prospective studies have noted around half of the patients with LC meet the diagnostic criteria for ME/CFS^9^, and roughly 75% of ME/CFS patients report an infection preceding symptom onset^6^.

Despite these similarities in clinical symptoms, there is no known molecular basis of ME/CFS and LC, with few available molecular signatures that explain the shared symptoms. Due to the lack of available molecular markers, there are neither standard diagnostic tests nor treatments for LC and ME/CFS. With no available options, patients with ME/CFS and/or LC can only be diagnosed with a physical and mental examination, based on the presentation of symptoms such as persistent fatigue and post-exertional malaise^7, 17^. With the mounting public health burden of ME/CFS and LC, there is an urgent need to understand the biochemical underpinnings of these conditions, which can guide the development of new diagnostics and therapies^6^.

Here we identify shared molecular signatures between ME/CFS and LC donors versus healthy controls, specifically signs of oxidative stress. We focused on immune cell bioenergetics based on several lines of evidence. Multiple studies^10-13^ have found signs of inflammation in LC and CFS patients, including higher serum cytokine levels^10-12^ that are associated with patient-reported fatigue severity^10^ and changes in both lymphocyte and monocyte cell populations^12^ among LC donors. These studies have suggested that the immune system may play a role in ME/CFS/LC pathogenesis and clinical symptoms.

Fatigue is the principal hallmark of ME/CFS and also one of the most common symptoms among LC^2,5,9^ patients. Since fatigue implies over and/or misallocation of cellular energy on disease processes, studying metabolism can meaningfully identify aberrations in how cells use and produce energy. Based on plasma metabolomic and proteomic studies, multiple metabolic pathway aberrations have been detected in LC and ME/CFS samples, including changes in serotonin and tryptophan metabolism^14^, cholesterol metabolism^51^, dysfunctional gut microbial butyrate biosynthesis pathways^15,16^, oxygen transport to tissues^72^, and deficient mitochondrial fatty acid oxidation and ATP metabolism^17, 20^.

We also focused on immune cells because they are a critical consumer of host energy and regulator of host metabolism. Specifically, the immune system accounts for 15-20% daily energy expenditure in humans^18,19^. This usage is thought to increase to 25%, during a serious infection, highlighting the significant energy expenditure required for maintenance, activation, and proliferation of immune cells^13-14^. Among the intracellular pathways in immune cells, mitochondrial metabolism is especially relevant. In addition to its central role in generating over 95% of a cell’s energy through ATP, changes in mitochondrial morphology and function accompany lymphocyte differentiation and activation^21^, with mitochondria driving cytokine production^22^, thereby linking energy metabolic deficits in immune cells with chronic inflammation. Along these lines, studies have hypothesized and identified signatures of mitochondrial dysfunction among CFS and LC patients, including down-regulation of host mitochondrial genes even after COVID-19 recovery^23^, altered mitochondrial morphology^24^ in ME/CFS donor T cells, decreases in mitochondrial membrane potential^25^ among recovered COVID-19 subjects’ lymphocytes, and redox dysregulation in both ME/CFS and COVID-19^26^.

Reactive oxygen species (ROS) are at the nexus of chronic inflammation and metabolic regulation, with its critical role in driving mitochondrial oxidative phosphorylation, inflammatory cytokine activation^28^, and tissue damage (characteristics of COVID-19 recovery)^28^. Based on the separate lines of evidence implicating chronic inflammation and metabolic dysregulation in ME/CFS and LC, we broadly profiled mitochondrial bioenergetic changes, specifically redox parameters, in lymphocytes from peripheral blood mononuclear cells (PBMCs) of 16 healthy controls, 15 ME/CFS, and 15 LC donors. From several measurements capturing ROS levels, oxidative damage, and changes in mitochondrial redox pathways, our findings identify elevated oxidative stress among lymphocytes is a shared molecular feature of ME/CFS and LC, associated with specific functional proliferation defects. Particularly striking was that only female patients had elevated ROS levels in ME/CFS and LC, together with the hyperproliferation of lymphocytes after stimulation in culture, whereas both sexes showed evidence of elevated levels of reduced glutathione and lipid oxidative damage. Thus, while there are major phenotypic differences between the sexes, they converge on evidence of oxidative stress and mitochondrial damage, which may lead to this type of immune system dysfunction acting as an “energy sink” analogous to a severe infection, leading to the persistent fatigue and other symptoms characteristic of these diseases.

## Results

### 1. Elevated reactive oxygen species in ME/CFS and Long COVID donor lymphocytes, compared to healthy controls

To profile bioenergetic parameters, we obtained healthy control, ME/CFS, and LC PBMCs. ME/CFS and LC donor PBMCs from the ME/CFS Collaborative Research Center and the Stanford Post-Acute COVID Syndrome clinic, respectively. Healthy control patient PBMCs were obtained from the Stanford Blood Bank, where donors were screened using a medical history questionnaire^30^. ME/CFS patients, including patients meeting the National Academy of Medicine (NAM) ME/CFS criteria before and after the start of the COVID-19 pandemic, were diagnosed by a physician using the National Academy of Medicine^7^ and Canadian Consensus criteria^29^. LC donors were diagnosed using the combination of the Center for Disease Control^17^ criteria for “Long COVID or post-COVID conditions” and a symptom and functional status questionnaire. Together, these measures capture whether patients report new symptoms four or more weeks after COVID infection and incorporates the Post-COVID 19 Functional Status (PCFS) Scale^27^. Additional details related to patient characteristics in each group are included in the methods section. Figure 1A shows that the mean ages are balanced across all three cohorts, and the proportion of females is roughly half among healthy controls and LC donors, with over 75% of ME/CFS donors being female. This distribution reflects the higher incidence of ME/CFS in women compared to men^31^.

**Figure 1.**
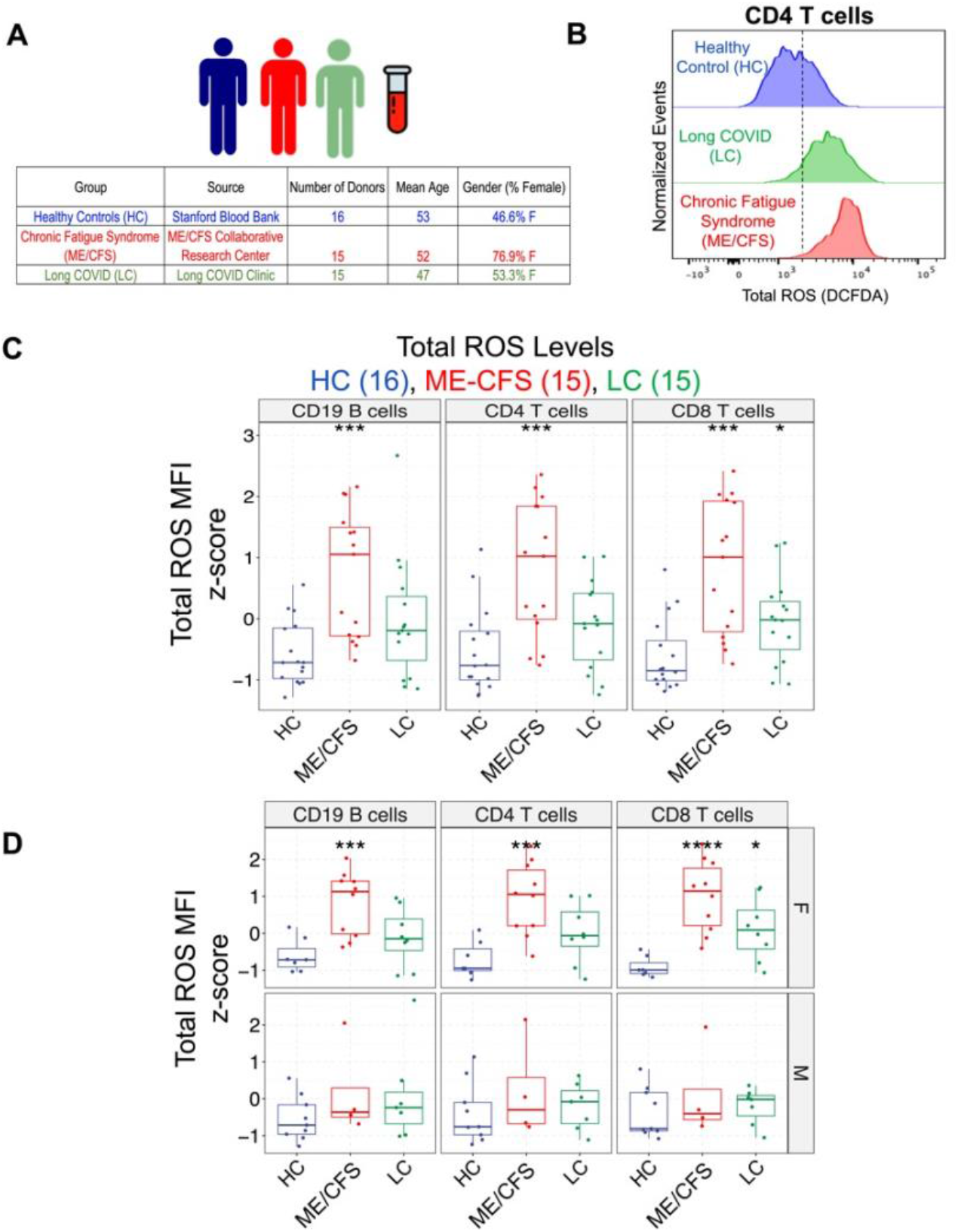
Comparison of total reactive oxygen species (ROS) levels in lymphocytes of healthy controls (HC), myalgic encephalomyelitis-chronic fatigue syndrome (ME/CFS), and Long COVID (LC) donors. (A) Peripheral blood mononuclear cells were characterized from 16 healthy control (HC), 15 Long COVID (LC), and 15 Myalgic Encephalomyelitis/Chronic Fatigue Syndrome (ME/CFS) samples. Age distributions were similar between all groups with gender distributions similar between HC and LC groups. (B) Representative flow cytometry plots for reactive oxygen species (ROS) indicator 2’,7’-dichlorodihydrofluorescein diacetate (DCFDA) are shown for CD4 T cells for a single healthy control (HC), Long COVID (LC), and ME/CFS donor. Y-axis represents normalized events to the mode. For C-E, * indicates p < 0.05, ** indicates p < 0.01, *** indicates p < 0.001 based on a two-sided t-test (C) Comparing 16 HC, 15 LC, and 15 ME/CFS donor lymphocytes, DCFDA staining shows higher total median fluorescence intensity (MFI) ROS levels in ME/CFS (CD4T p = 5.60*10^-4^, CD8T p = 1.91*10^-4^, CD19B p = 3.04*10^-4^) and LC (CD4 T cells p = 0.104, CD8 T cells p = 0.0210, CD19 B cells p = 0.0727) lymphocytes compared to healthy controls. (D) Comparison of MFI ROS levels uniquely identifies higher ROS levels in ME/CFS and LC females, compared to controls (females ME/CFS v. HC: CD19 B cells p = 0.00065, CD4 T cells p = 0.00045, CD8 T cells p = 9.5*10^-5^; LC v. HC: CD19 B cells p = 0.144, CD4 T cells p = 0.062, CD8 T cells p = 0.01065). In contrast, ME/CFS and LC males show no significant difference (males ME/CFS v. HC: CD19 B cells p = 0.357, CD4 T cells p = 0.469, CD8 T cells p = 0.516; males ME/CFS v. LC: CD19 B cells p = 0.278, CD4 T cells p = 0.643, CD8 T cells p = 0.583)

Using flow cytometry, we compared mitochondrial bioenergetic parameters in CD19 B cells as well as CD4 and CD8 T cell lymphocyte populations. Sample gating strategy for identifying these lymphocyte populations is shown in Supplementary Figure S1A. Although we did not identify any differences between healthy controls and ME/CFS or LC donors in mitochondrial mass or membrane potential (Supplementary Figure S2), decreases in mitochondrial ATP levels were observed across ME/CFS (1.6x decrease on average; HC v. ME/CFS two-sided t-test CD4 T cells p = 0.0458, CD8 T cells p = 0.0974, CD19 B cells p = 0.356) and LC donors (2.8x decrease on average; HC v. LC two-sided t-test CD4 T cells p = 0.0076, CD8 T cells p = 0.0182, CD19 B cells p = 0.0842), compared to healthy controls (Supplementary Figure S2). Our findings are consistent with a published study showing oxidative phosphorylation defects among PBMCs in CFS patients, compared to healthy controls^69^. As the decrease in mitochondrial ATP levels implies functional mitochondrial deficits and mitochondria significantly contribute to cellular ROS production through oxidative phosphorylation, we measured total ROS levels. Flow cytometric staining using 2’,7’ –dichlorofluorescein diacetate (DCFDA) identified elevated reactive oxygen species (ROS) levels among lymphocytes in ME/CFS and LC donors, compared to healthy controls (Figure 1B, 1C). DCFDA is hydrolyzed intracellularly and oxidized by ROS, including hydroxyl radicals, hydrogen peroxide derived compounds, to form 2′,7′-dichlorofluorescein (DCF). The formation of DCF is associated with increased fluorescence, which can be detected through flow cytometry. Representative flow cytometric plots from a single HC, LC, and ME/CFS donor for CD4 T cells are shown in Fig. 1B. Comparisons of DCF median fluorescence intensity (MFI) measurements between groups, corresponding to ROS levels, indicate statistically significant increases in both LC (n=15) and ME/CFS (n=15) donor lymphocytes, compared to healthy controls (n=16) (Figures 1C) (ME/CFS CD4T p = 5.60*10^-4^, CD8T p = 1.91*10^-4^, CD19B p = 3.04*10^-4^; LC CD4 T cells p = 0.104, CD8 T cells p = 0.021, CD19 B cells p = 0.073).

Closer examination of the distribution of MFI ROS values among ME/CFS donors (Figure 1C, 1D) revealed a bimodal distribution, with 8/15 donors (Fig. 1C) exhibiting over 2-4x higher ROS levels compared to healthy controls. We found this bimodal distribution could be explained by patient sex, where total ROS levels appear to be uniquely elevated in female ME/CFS and LC donors (Figure 1D). Among females (top row, Figure 1D), total ROS levels are significantly elevated in ME/CFS and LC donors, compared to controls (females ME/CFS v. HC: CD19 B cells p = 0.00065, CD4 T cells p = 0.00045, CD8 T cells p = 9.5*10^-5^; LC v. HC: CD19 B cells p = 0.144, CD4 T cells p = 0.062, CD8 T cells p = 0.01065). No differences in ROS levels are found between males with ME/CFS or LC, compared to control males (Figure 1E, bottom row) (males ME/CFS v. HC: CD19 B cells p = 0.357, CD4 T cells p = 0.469, CD8 T cells p = 0.516; males ME/CFS v. LC: CD19 B cells p = 0.278, CD4 T cells p = 0.643, CD8 T cells p = 0.583). Notably, our data offers one potential way to interpret several epidemiological studies^31-34^ that compare post-COVID 19 and ME/CFS symptoms incidence between males and females. These studies estimate that ME/CFS is three to four times more common in women than men^31^, with significantly higher likelihood for females to develop post COVID-19 symptoms^32-34^ (e.g., 3.3x odds ratio estimate for LC in females v. males in Ref. 32).

Additionally, The Post-COVID 19 Functional Status (PCFS) Scale^18^ and Bell disability score^9^ were used to assess fatigue severity. These patient-reported fatigue measures capture a patient’s functional status, with a fully functional patient having no symptoms at rest and able to work full-time and an extremely sick patient unable to work, bedridden, and experiencing severe fatigue symptoms on a continuous basis. We did not detect any association between LC/CFS patient-reported fatigue severity and total ROS levels (Supplementary Figure S3A), irrespective of gender.

Since the duration of symptoms could be pinpointed for LC patients, we also evaluated whether DCF MFI levels in LC patients are associated with duration of LC symptoms. Although we did not detect a significant association (Supplementary Figure S3B), we found that female LC subjects consistently showed a weak positive association and males showed a weak negative association between symptom duration and total ROS levels across lymphocyte populations (females CD19B R = 0.59 p = 0.12, CD4T R = 0.49 p = 0.22, CD8T R = 0.5 p = 0.21; males CD19B R = -0.91 p = 0.0039, CD4T R = -0.63 p = 0.13, CD8T R = -0.43 p = 0.34). These divergent trends in ROS levels suggests that the LC and ME/CFS disease pathogenesis is also distinct between males and females, specifically the mechanisms underlying resolution of sustained oxidative stress.

While we observed no significant association between ROS levels and age (Supplementary Figure S3C), we did identify a positive correlation among LC donors between BMI and total ROS levels (CD19 B cells R = 0.56, p = 0.028; CD4 T cells R = 0.46, p = 0.081; R = 0.14, p = 0.13) (Supplementary Figure S3D). As BMI has also been positively correlated with the incidence of post-COVID 19 symptoms^34^, our findings suggest that differences in oxidative stress tracks with clinical heterogeneity in ME/CFS and LC. Moreover, these results highlight that elevated ROS levels in lymphocytes are a more sensitive molecular marker of ME/CFS and LC in females, particularly those with higher BMI.

### 2. Changes in ROS levels are accompanied by alterations in oxidative stress pathways

Based on our results showing differences in ROS levels, we specifically investigated how ROS pathways are altered in ME/CFS and LC donors, including those related to anti-oxidant clearance and subsequent oxidative damage. Figure 2A summarizes some of the critical ROS generation and clearance pathways^35,36^, for which we conducted several measurements to evaluate differences between HC and LC/ME/CFS groups. As depicted in Figure 2A, intracellular ROS levels are mediated by the balance between ROS-generation and antioxidant processes. In terms of ROS-generation pathways, calcium can drive the production of mitochondrial ROS through electron transport chain activity coupled to NADPH oxidases. To regulate mitochondrial ROS levels (e.g., superoxide (O_2_^-^)), superoxide dismutase 2 (SOD2) converts superoxide to hydrogen peroxide (H_2_O_2_). Hydrogen peroxide can be subsequently transformed to H_2_O and O_2_ through catalase and glutathione peroxidase, which is coupled to oxidation of glutathione (GSH). Fig. 2A also summarizes the measured sex-specific differences in ROS generation, clearance, and down-stream pathways, where ME/CFS females show elevated mitochondrial calcium (Fig. 2B) and ROS levels (Fig. 1C), whereas ME/CFS males show larger changes in lipid oxidative damage (Fig. 3F), compared to healthy counterparts.

**Figure 2.**
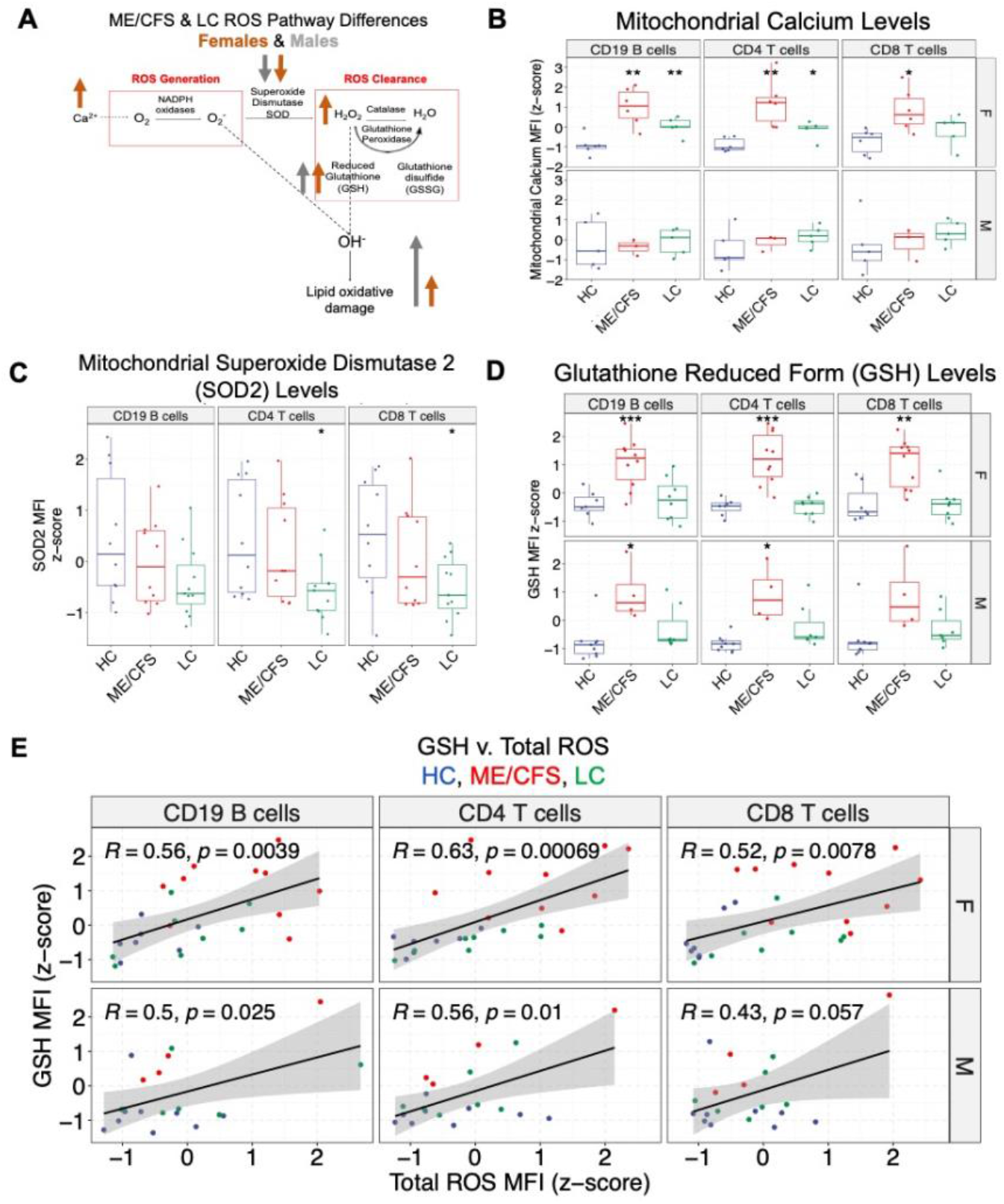
Comparison of important intermediates and proteins involved in ROS generation and clearance within lymphocytes between HC, LC, and ME/CFS donors. (A) Summary of ROS generation and clearance pathways, including mitochondrial ROS (e.g., superoxide O_2-_) calcium and NADPH oxidase-mediated generation, superoxide dismutase mediated-conversion to hydrogen peroxide, and hydrogen peroxide breakdown by catalase and glutathione. The alterations in these ROS generation and clearance pathways among ME/CFS, LC donors are shown by sex (females shown in brown, males in gray). Upward arrow corresponds to elevation of parameter in ME/CFS and LC donors, compared to healthy controls of the same gender. The rest of Fig. 2 and Fig. 3 includes the data supporting these sex-specific differences. *For B-D,* * indicates p < 0.05, ** indicates p < 0.01, *** indicates p < 0.001 based on a two-sided t-test (B) Comparison of mitochondrial calcium levels based on Rhod-2 AM flow cytometric staining identifies statistically significant higher levels in ME/CFS and LC female donors (females: ME/CFS v. HC CD19B p = 0.00238, CD4T p = 0.0063, CD8T p = 0.0137; ME/CFS v. LC CD19B p = 0.0097, CD4T p = 0.0172, CD8T p = 0.240; males: ME/CFS v. HC CD19B p = 0.791, CD4T p = 0.514, CD8T p = 0.825; LC v. HC CD19B p = 0.851, CD4T p = 0.241, CD8T p = 0.358). (C) Flow cytometric comparison of mitochondrial superoxide dismutase 2 (SOD2) protein levels finds significantly lower SOD2 levels in LC donors, compared to healthy controls (HC v. ME/CFS CD19 B cells p = 0.326, CD4 T cells p = 0.638, CD8 T cells p = 0.384; HC v. LC CD19 B cells p = 0.073, CD4 T cells p = 0.023, CD8 T cells p = 0.023). (D) Comparison of glutathione (GSH) levels finds significantly higher MFI levels in both male and female ME/CFS donors (females ME/CFS v. HC CD19 B cells p = 0.00048, CD4 T cells p = 0.00014, CD8 T cells p = 0.00159; LC v. HC CD19B cells p = 0.580, CD4 T cells p = 0.827, CD8T cells p = 0.958; males ME/CFS v. HC CD19 B cells p = 0.0327, CD4 T cells p = 0.0354, CD8 T cells p = 0.0894; LC v. HC CD19 B cells p = 0.196, CD4 T cells p = 0.093, CD8 T cells p = 0.278). (E) Based on the reaction between hydrogen peroxide (H_2_O_2_) and glutathione (GSH) in Figure 2A (H_2_O_2_ + 2GSH ßè2H_2_O + GSSG), the association between GSH and total ROS levels (Fig. 1C) is shown for both females (top row) and males (bottom row). With each point corresponding to a donor, the plot shows a statistically significant *positive* correlation between total ROS and GSH levels (females CD19 B cells R = 0.56, p = 0.0039, CD4 T cells R = 0.63 p = 6.9*10^-4^; CD8 T cells R = 0.52 p = 7.8*10^-3^; males CD19 B cells R = 0.5 p = 0.025, CD4 T cells R = 0.56 p = 0.01, CD8 T cells R = 0.43 p = 0.057).

**Figure 3.**
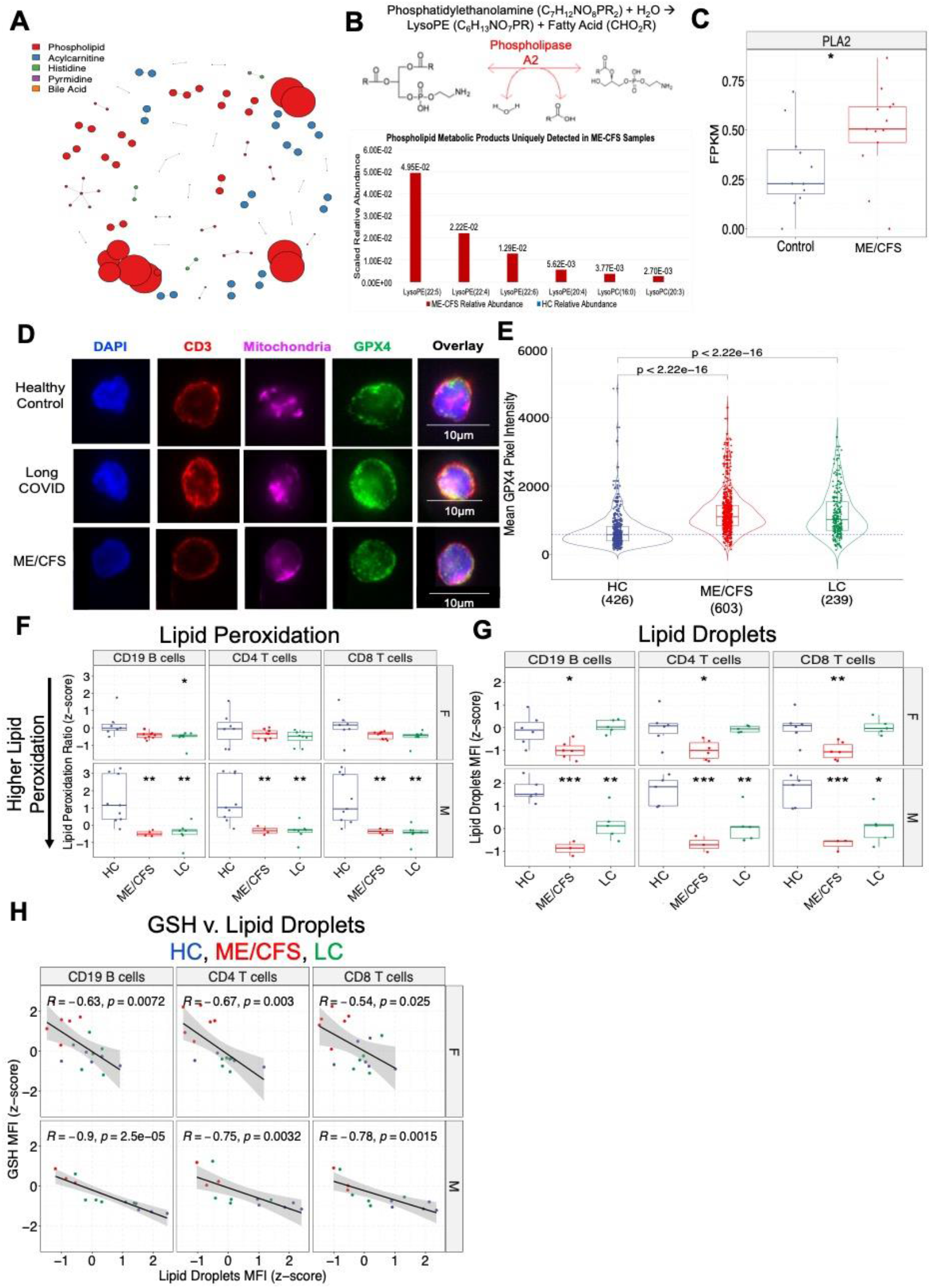
Elevated lipid oxidative damage in ME/CFS and LC donor lymphocytes, compared to healthy controls. (A) Using extracted metabolites from 200,000 sorted CD3 T cells from ME/CFS and HC donors, hydrophilic interaction liquid chromatography-mass spectrometry and Turium analysis were conducted to map metabolic pathway differences between ME/CFS donor and healthy control T cells. From 747 identified metabolites, the top five mapped pathway differences are shown. Size of the nodes correspond to magnitude of the difference in relative abundance of detected metabolites from mass spectrometry between ME/CFS and HC samples. (B) The reactions encoded by the large red circles (Figure 3A) are shown, corresponding to reactions significantly up-regulated in CFS donors compared to healthy controls. The phospholipid synthesis reactions involve the production of lysoPE metabolites and are catalyzed by phospholipase A2 (PLA2). (C) Comparison of PLA2 transcript levels from bulk RNA-seq data (Bouquet et al., Ref. 42) between 11 sedentary controls and 14 CFS patients identifies significantly higher levels of PLA2 in CFS donors (p < 0.05, two-sided t-test). Transcript levels are compared using fragments per kilobase of transcript per million mapped reads (FPKM). (D) Representative images from immunofluorescence staining are shown for glutathione peroxidase 4 (GPX4) in CD3 T cells from 1 HC, 1 LC, and 1 ME/CFS donor (blue corresponds to DAPI, red to CD3, magenta to MitoTracker Far Red/mitochondria, green to GPX4) (E) Quantification of average GPX4 pixel intensity from 1 HC, 1 LC, and 1 ME/CFS donor. Each point corresponds to a CD3 T cell, with comparisons conducted across 426 HC, 603 ME/CFS, and 239 LC T cells from maximum projection images. Significant increases in average pixel intensity indicate higher GPX4 levels in ME/CFS and LC donors, compared to healthy controls (p < 2.2*10^-16^ for both ME/CFS and LC from two-sided t-test). (F) Flow cytometric comparison of lipid peroxide levels, using ratiometric lipid peroxidation sensor, shows both ME/CFS and LC donor lymphocytes have significantly higher lipid peroxidation, compared to healthy controls. Due to lower lipid peroxide levels in healthy controls, male ME/CFS and LC donors exhibit larger changes compared to females (males ME/CFS v. HC: CD19 B cells p = 0.0027, CD4 T cells p = 0.0037, CD8 T cells p = 0.0041, LC v. HC CD19 B cells p = 0.003, CD4 T cells p = 0.0033, CD8 T cells p = 0.0029; females ME/CFS v. HC: CD19 B cells p = 0.076, CD4 T cells p = 0.448, CD8 T cells p = 0.131, LC v. HC: CD19 B cells p = 0.0365, CD4 T cells p = 0.236, CD8 T cells p = 0.0977). Lower y-axis values correspond to *higher* lipid peroxidation. (G) Flow cytometric comparison of lipid droplet levels identifies lower lipid droplet levels in ME/CFS donors compared to HC across both males and females (males ME/CFS v. HC: CD19 B cells p = 1.52*10^-4^, CD4 T cells p = 5.92*10^-4^, CD8 T cells p = 7.14*10^-4^, LC v. HC: CD19 B cells p = 7.31*10^-3^, CD4 T cells p = 9.39*10^-3^, CD8 T cells p = 0.0114; females ME/CFS v. HC CD19 B cells p = 0.0242, CD4 T cells p = 0.0205, CD8 T cells p = 0.00876; LC v. HC CD19 B cells p = 0.622, CD4 T cells p = 0.826, CD8 T cells p = 0.974). (H) Negative association between lipid droplet (Figure 3G) and GSH levels (Figure 2D) among lymphocytes (females: CD19 B cells R = -0.63, p = 0.0072, CD4 T cells R = -0.67 p = 0.003, CD8 T cells R = -0.54 p = 0.025; males: CD19 B cells R = -0.9, p = 2.5*10^-5^, CD4 T cells R = -0.75 p = 0.0032, CD8 T cells R = -0.78, p = 0.0015).

From flow cytometry analysis, we compared mitochondrial Ca^2+^ levels using Rhod-2 AM, a high affinity calcium indicator that primarily localizes in mitochondria. Compared to healthy controls, lymphocytes from ME/CFS and LC donors have 1.67x and 1.32x respectively higher calcium levels, with CD4 T cells showing statistically significant elevations in ME/CFS and LC groups compared to healthy controls (ME/CFS CD19 p = 0.014, CD4 p = 0.003, CD8 p = 0.03; LC CD19 p = 0.221, CD4 p = 0.023, CD8 = 0.132) (Supplementary Figure 4A). Notably, the mitochondrial calcium levels were uniquely elevated in female ME/CFS and LC donor lymphocytes, compared to healthy controls (Figure 2B) (females: ME/CFS v. HC CD19B p = 0.00238, CD4T p = 0.0063, CD8T p = 0.0137; LC v. HC CD19B p = 0.0097, CD4T p = 0.0172, CD8T p = 0.240; males: ME/CFS v. HC CD19B p = 0.791, CD4T p = 0.514, CD8T p = 0.825; LC v. HC CD19B p = 0.851, CD4T p = 0.241, CD8T p = 0.358). We also compared SOD2 levels (Figure 2C) between healthy controls and ME/CFS and LC subjects. Our comparison found that LC donor CD4 and CD8 T cells have significantly lower SOD2 levels, compared to healthy controls (ME/CFS CD19 p = 0.326, CD4 p = 0.63, CD8 p = 0.38; LC CD19 p = 0.073, CD4 p = 0.023, CD8 p = 0.023). These differences in SOD2 levels appeared to be similar across both males and females (Supplementary Figure 4B).

Combining these differences to capture the balance between mitochondrial ROS drivers and anti-oxidant pathways, we find the ratio of Ca^2+^ to SOD2 MFI levels is highly elevated in *both* the ME/CFS and LC donors, compared to healthy controls. The mitochondrial calcium to SOD2 ratio is elevated by 1.67x in LC donors and 1.75x in ME/CFS donors across lymphocytes, compared to healthy controls (ME/CFS: CD19 p = 0.011, CD4 p = 0.005, CD8 p = 0.025; LC: CD19 p = 0.05, CD4 p = 0.006, CD8 p = 0.025). Importantly, these results further show that mitochondrial dysfunction is shared between ME/CFS and LC donors.

To avoid cellular oxidative damage, glutathione can act as a ROS scavenger, where the reduced form of glutathione (GSH) reduces H_2_O_2_ to restore cellular redox balance ^26, 37^. Glutathione acts as an important reducing agent in several cellular redox reactions, including those catalyzed by glutathione reductase, glutathione peroxidase, catalase. The redox balancing reaction between hydrogen peroxide and glutathione is shown in Figure 2A and can be written as follows: H_2_O_2_ + 2GSH ←→ 2H_2_O + GSSG. Based on this reaction and the hypothesis that endogenous glutathione deficiency is a driver of severe COVID-19 disease^37^, we compared GSH levels using flow cytometry between HC, LC, and ME/CFS donors. Surprisingly, our comparisons found significantly higher GSH levels in ME/CFS donors, across all lymphocyte populations (HC v. ME/CFS CD19 B cells p = 2.0*10^-6^, CD4 T cells p = 5.7*10^-7^, CD8 T cells p = 1.7*10^-5^; HC v. LC CD19 B cells p = 0.14, CD4 T cells p = 0.064, CD8 T cells p = 0.38). As we identified higher ROS levels *only* in females (Figure 1D), we also compared glutathione levels in ME/CFS and LC males and females. Our comparison (Figure 2D) found both ME/CFS males and females exhibit higher GSH levels, compared to controls of the same gender (females ME/CFS v. HC CD19 B cells p = 0.00048, CD4 T cells p = 0.00014, CD8 T cells p = 0.00159; LC v. HC CD19B cells p = 0.580, CD4 T cells p = 0.827, CD8T cells p = 0.958; males ME/CFS v. HC CD19 B cells p = 0.0327, CD4 T cells p = 0.0354, CD8 T cells p = 0.0894; LC v. HC CD19 B cells p = 0.196, CD4 T cells p = 0.093, CD8 T cells p = 0.278). Since DCFDA inference of total ROS (Figure 1C) can capture cellular hydrogen peroxide levels^38^, we also evaluated the association between total ROS and GSH measurements (Figure 2E). As shown in Figure 2E, we noted a significant positive correlation between total ROS and GSH levels across all lymphocyte populations (females CD19 B cells R = 0.56, p = 0.0039, CD4 T cells R = 0.63 p = 6.9*10^-4^; CD8 T cells R = 0.52 p = 7.8*10^-3^; males CD19 B cells R = 0.5 p = 0.025, CD4 T cells R = 0.56 p = 0.01, CD8 T cells R = 0.43 p = 0.057). Importantly, although ME/CFS and LC female lymphocytes uniquely exhibit higher total ROS levels, higher GSH levels across both males and females highlight that response to oxidative stress is a common feature in ME/CFS and LC lymphocytes across both sexes.

### 3. Elevated oxidative stress in ME/CFS and LC donors is associated with lipid damage

We also probed the likely down-stream metabolic consequences of oxidative stress, specifically whether there was evidence of oxidative damage in known pathways^39^.

To study differences in these pathways, we separately pooled 200,000 sorted CD3^+^ T cells from both healthy controls and ME/CFS donors and immediately extracted intracellular metabolites. Using hydrophilic interaction liquid chromatography mass spectrometry^40^, 45,507 unique *m/z* analytes were detected in samples, of which 747 metabolites were identified using analytical standards. From mass spectrometry data, systems chemistry analysis was conducted using Turium^24^, a computational program which enabled the mapping of 747 metabolites to 327 pathways. This approach helped identify the top pathways that are increased or uniquely detected in ME/CFS patients, compared to healthy controls (Figure 3A). In Figure 3A, each circle corresponds to a specific reaction, up-regulated in ME/CFS donor T cells, compared to healthy controls. The size of the node corresponds to the magnitude of the relative abundance difference between ME/CFS and healthy control samples for the detected metabolite from the mass spectrometry data. We found several metabolic pathway differences, with the most significant corresponding to phospholipid synthesis (Figure 3A). Specifically, we found that phospholipid metabolites were significantly elevated in ME/CFS samples compared to healthy controls, including lysophosphatidylethanolamine (lysoPE) products (lysoPE(22:5), lysoPE(22:4), lysoPE(22:6), lysoPE(20:4), lysoPE(16:0), lysoPE(20:3)) (Figure 3B). The reaction to produce these lysoPE products is shown in Figure 3B, (Phosphotidylethanolamine (C_7_H_12_NO_8_PR_2_) + H_2_O → LysoPE(C_6_H_13_NO_7_PR) + CHO_2_R). These results are consistent with earlier plasma metabolomic studies on ME/CFS that identified significant phospholipid abnormalities^17^.

As the identified reaction products (Figure 3B) are catalyzed by phospholipase A2 (PLA2), we used RNA-seq data from Bouquet et al.^42^ to test for differences in PLA2 transcript expression from 11 sedentary controls and 14 CFS patients whole blood samples. While Bouquet et al. studied the effects of cardiopulmonary exercise on CFS patients, we specifically compared the phospholipase A2 levels (Figure 3C) at the 7 days post-exercise timepoint, when CFS symptoms returned in patients^42^ and lymphocytes were least likely to reflect the effects of exercise. We found that PLA2G4A, a calcium dependent cystolic phospholipase implicated with eicosanoid metabolism, has significantly higher transcript levels in ME/CFS donors compared to controls (p = 0.044, two-sided t-test) at day 7 of this study (Figure 3C). These findings provide an external validation for phospholipid synthesis dysregulation in ME/CFS donors.

Lipid peroxidation of fatty acyl groups occurs predominantly in membrane phospholipids, where lipid peroxidation promotes the formation of several lipid byproducts including lysophospholipids^43^. Based on the differences in phospholipid synthesis and changes in glutathione levels, we evaluated whether ME/CFS and LC donors exhibited signs of lipid oxidative damage, specifically lipid peroxidation.

The conversion of lipid peroxides to lipid alcohols is catalyzed by glutathione peroxidase 4 (GPX4), which protects lipids from ROS damage. Therefore, we conducted immunofluorescence staining of GPX4 (Figure 3D), across CD3 T cells from a single HC, LC, and ME/CFS donor. Comparison of GPX4 mean pixel intensity from immunofluorescence staining across 426 HC, 603 ME/CFS, and 239 LC CD3 T cells shows that LC and ME/CFS donor lymphocytes have higher (1.77x and 1.9x, respectively) GPX4 levels (Figure 3E), when compared to T cells from healthy controls (p < 2.22 * 10^-16^ for both ME/CFS and LC v. HC comparisons, two-sided t-test). Next, using flow cytometry, we measured lipid peroxide levels in HC, ME/CFS, and LC donor lymphocytes, using a ratiometric lipid peroxidation sensor. Our measurements found substantially higher lipid peroxide levels across all lymphocytes in both ME/CFS and LC donors, compared to healthy controls (Supplementary Figure 4C), lower y-axis values correspond to higher lipid peroxidation) (HC v. ME/CFS CD19 B cells p = 0.000995, CD4 T cells p = 0.00556, CD8 T cells p = 0.00155; HC v. LC CD19 B cells p = 0.000528, CD4 T cells p = 0.00281, CD8 T cells p = 0.000988). Both ME/CFS and LC males and females presented with similar levels of lipid peroxides (Figure 3F). However, the differences between controls and ME/CFS/LC donors in lipid peroxide levels were significantly more pronounced in males, due to lower levels of lipid peroxides among males compared to female controls (males ME/CFS v. HC: CD19 B cells p = 1.52*10-4, CD4 T cells p = 5.92*10-4, CD8 T cells p = 7.14*10-4, LC v. HC: CD19 B cells p = 7.31*10-3, CD4 T cells p = 9.39*10-3, CD8 T cells p = 0.0114; females ME/CFS v. HC CD19 B cells p = 0.0242, CD4 T cells p = 0.0205, CD8 T cells p = 0.00876; LC v. HC CD19 B cells p = 0.622, CD4 T cells p = 0.826, CD8 T cells p = 0.974).

To protect membranes from lipid oxidative damage, the formation of lipid droplets can help restore redox homeostasis^26^ by sequestering ROS damaged lipids. Additionally, fatty acids derived from lipid droplets can be converted to acylcarnitines, the substrates for mitochondrial fatty acid oxidation. Based on our results in Figure 3A, which also detected changes in acylcarnitine metabolism in ME/CFS donors, we compared lipid droplet levels using HCS LipidTox Green Neutral Lipid staining, which stains for neutral intracellular lipid droplets. As shown in Supplementary Figure 4G, we found lower lipid droplet levels in both LC and ME/CFS donors across all measured lymphocyte populations (ME/CFS: CD19 p = 5.19*10^-4^, CD4 p = 4.58*10^-4^, CD8 p = 3.01*10^-4^; LC: CD19 p = 0.140, CD4 p = 0.0669, CD8 p = 0.0759). Both ME/CFS and LC males and females presented with decreases in lipid droplet levels (Figure 4G), although lipid droplet levels were higher in control females compared to males (males ME/CFS v. HC: CD19 B cells p = 1.52*10-4, CD4 T cells p = 5.92*10-4, CD8 T cells p = 7.14*10-4, LC v. HC: CD19 B cells p = 7.31*10-3, CD4 T cells p = 9.39*10-3, CD8 T cells p = 0.0114; females ME/CFS v. HC CD19 B cells p = 0.0242, CD4 T cells p = 0.0205, CD8 T cells p = 0.00876; LC v. HC CD19 B cells p = 0.622, CD4 T cells p = 0.826, CD8 T cells p = 0.974). Furthermore, across both males and females, lipid droplet levels are inversely correlated with GSH levels across B and T lymphocytes. Figure 3H shows the correlation across males and females (females: CD19 B cells R = -0.63, p = 0.0072, CD4 T cells R = -0.67 p = 0.003, CD8 T cells R = -0.54 p = 0.025; males: CD19 B cells R = -0.9, p = 2.5*10-5, CD4 T cells R = -0.75 p = 0.0032, CD8 T cells R = -0.78, p = 0.0015).

**Figure 4.**
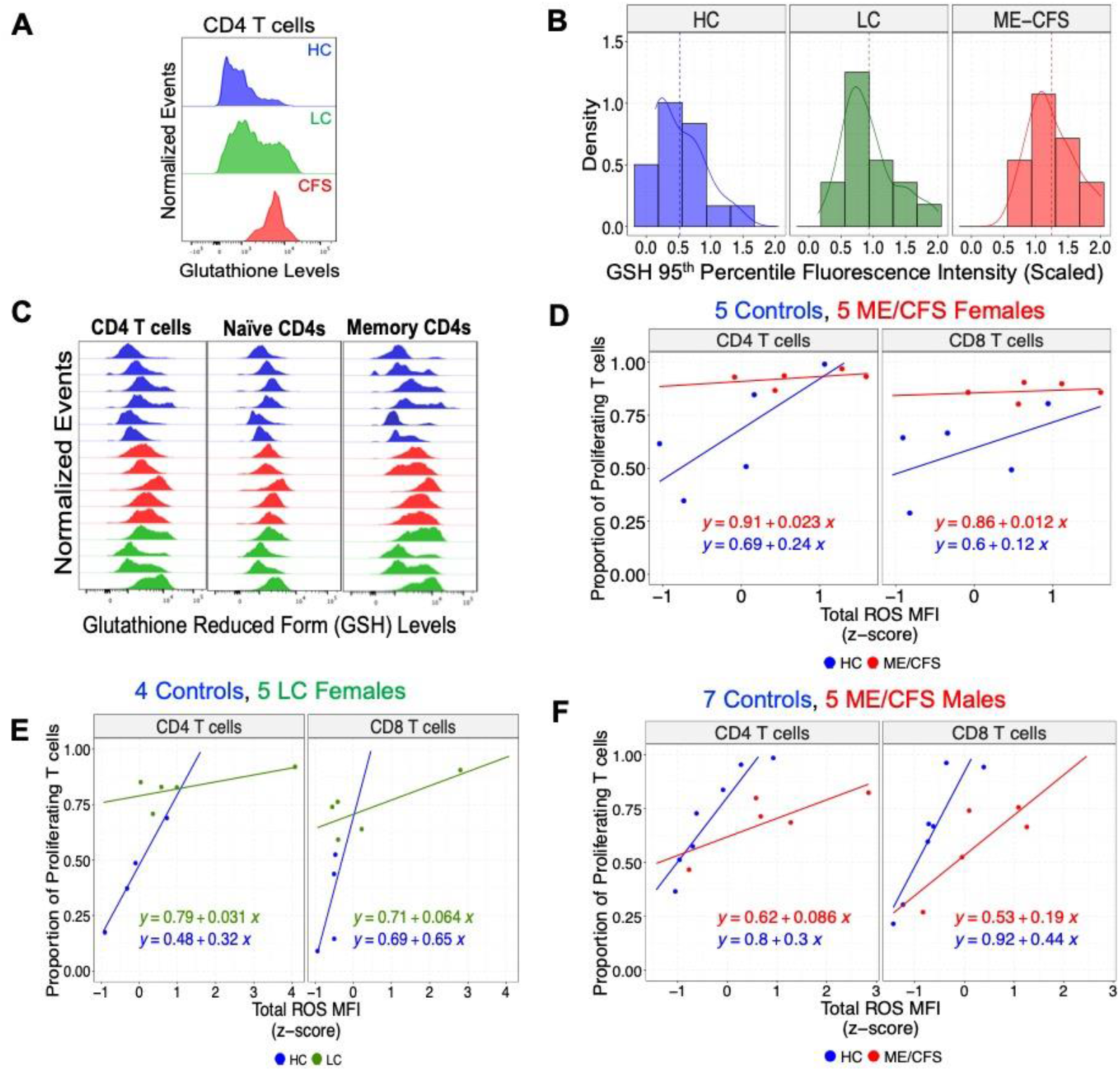
Effects of ROS on Adaptive Immune Response. (A) Representative glutathione (GSH) profile from flow cytometry staining for a HC, LC, and ME/CFS CD4 T cell shows a long-tailed distribution for both HC and LC donors and a shifted distribution for ME/CFS. (B) To study tail phenomena, the distribution of 95^th^ percentile from flow cytometry GSH data is plotted across HC, LC, and ME/CFS donors. Each group demonstrates distinct tail-decaying behavior, where healthy controls decay quickly (light-tailed distribution), LC show heavy-tailed decay, and ME/CFS show a shifted and bounded tail distribution. (C) To understand this decaying behavior at the tail of the distribution, we compared the GSH levels on an additional set of 5 HC, 4 LC, and 5 ME/CFS donors between total, naïve (CCR7^+^CD45RO^-^), and memory CD4 T cell populations (CD45RO^+^). The long tails are found *only* in memory and not naïve CD4 T cells, indicating that memory T cells are the main contributors to high oxidative stress in T cells. For D-F, to probe functional effects of oxidative stress, PBMCs were labeled with CellTraceViolet and stimulated with anti-CD3/CD28 beads and IL-2. Based on a dye dilution assay, the proportion of proliferating T cells as a function of total ROS levels are shown at 5 days post-stimulation. *Experiments for D-F were conducted separately*. (D) Data is shown for a set of 5 control and 5 ME/CFS females, for which female ME/CFS T cells hyperproliferate and exhibit higher ROS levels compared to control counterparts. The regression lines also show that control female CD4 and CD8 T cell proliferation has 10x greater slope with respect to ROS, compared to ME/CFS females. Compared to controls, the flat slope and higher line for ME/CFS T cells highlights hyperproliferation, suggesting a potential source of patient fatigue. (E) Consistent with Fig. 4D, data shown for 4 control and 5 LC females shows hyperproliferation among LC CD4 and CD8 T cells. The regression lines also show that control female CD4 and CD8 T cell proliferation has 10x greater slope with respect to ROS, compared to LC females. Compared to controls, the flat slope and higher line for LC T cells highlights hyperproliferation, consistent with the data in Fig. 4D. (F) Data shown for 7 control and 5 ME/CFS males shows a distinct pattern from females. CD4 and CD8 T cells from ME/CFS males do not hyperproliferate compared to controls, as ME/CFS males with similar ROS levels to controls do not show higher proliferation. As T cell proliferation among controls show 3.5x and 2.3x greater slope compared to ME/CFS CD4 and CD8 T cell counterparts, these results show greater insensitivity to ROS levels among ME/CFS male donors.

Our results correlating lipid droplet and glutathione levels link redox homeostasis with lipid composition and fatty acid oxidation, highlighting these metabolic consequences of oxidative stress. Furthermore, our findings highlight gender-specific redox differences that likely relate to distinct pathophysiology mechanisms among ME/CFS/LC males and females (Figure 2A).

### 4. Elevated oxidative stress in memory CD4T and altered T cell proliferation responses to ROS in ME/CFS donors suggests a deficient adaptive immune response, upon stimulation

Beyond the metabolic effects, we also probed the functional consequences of excess oxidative stress in ME/CFS donor lymphocytes. It is well known that elevated ROS levels are a critical, albeit transient component of both B and T cell lymphocyte activation^45, 46,74^. Specific to T cells, upon T cell activation, TCR signaling stimulates calcium influx into the mitochondria, which drives mitochondrial ROS production, activates NFAT signaling, and triggers IL2 cytokine production. Moreover, T cells without mitochondrial ROS are unable to proliferate upon antigen stimulation^47^. Thus, the constitutive ROS elevation and alterations in mitochondrial calcium levels (Figure 2B) that we see here in ME/CFS females suggests that there will be sex-specific differences in T cell proliferation compared to controls.

Next, re-examination of glutathione (GSH) data revealed significant variability in GSH distribution among CD4 T cells *within* each donor, especially among HC and LC patients (Figure 4A). Comparing the extreme value distributions (95^th^ percentile fluorescence intensity) between HC, LC, and ME/CFS donor CD4 T cells, we observed that the HC CD4s decay most quickly (light-tailed distribution), whereas LC CD4s decay with a heavy-tailed distribution (Figure 4B). In contrast, CFS donor CD4s have a shifted and bounded extreme value distribution. As comparisons of the extreme values alone *rather* than the medians or means identifies statistically significant elevation of GSH in LC and ME/CFS donors (HC v. LC CD4 p = 0.00831; HC v. ME/CFS CD4 p = 8.5*10^-6^), this result underscores that only comparing the medians overlooks the critical heterogeneity that contributes to LC heavy-tail behavior. These results also suggest that a significantly larger proportion of T cells endure oxidative stress in ME/CFS donors, which is reflected in the shift in both comparisons of medians (Figure 2D) and extreme value distributions (Fig. 4B). To assess which T cells may account for the differences in these phenomena, we compared naïve and memory CD4 T cell populations from 5 additional HC, 5 ME/CFS donors, and 4 LC subjects (Figure 4C). Using CCR7 and CD45RO+ surface markers to delineate memory CD4 T cell populations (Supplementary Figure 1B), our analysis found that the long tails in CD4 GSH profiles were *only* reflected in memory and not naïve CD4 T cells (Figure 4C). Moreover, ME/CFS and LC memory populations with a heavy tail have higher GSH values compared to their healthy counterparts. As antigen stimulation encourages the formation and recall of a T cell memory response, our results also suggested that ME/CFS donor T cells would likely respond distinctly to antigen stimulation.

Therefore, we investigated the relation between oxidative stress in ME/CFS donor T cells and its proliferation, after stimulation. In light of our total ROS and mitochondrial calcium findings, we studied T cell proliferation in 5 HC and 5 ME/CFS female donor PBMCs. These PBMCs, which were labeled with CellTrace Violet proliferation dyes, were stimulated with anti-CD3/anti-CD28 antibodies and IL-2. The extent of proliferating T cells was measured 5 days post-stimulation, along with levels of oxidative stress and surface activation marker levels (CD69, CD137) in T cells (see methods for details). Separate experiments were conducted for ME/CFS females (Fig. 4D), LC females (Fig. 4E), and ME/CFS males (Fig. 4F). Upon stimulation, we observed no differences in CD69 and CD137 activation status for any ME/CFS and LC samples compared to HCs (Supplementary Figure S5). However, we noted sex-specific differences in the relationship between ROS and T cell proliferation, as described further below.

Consistent with the vital role of ROS in T cell activation, our analysis found among controls that the proportion of proliferating T cells linearly scales with oxidative stress (Fig. 4D-F). In contrast, the relation between ROS and T cell proliferation appears distinct in ME/CFS and LC subjects, with additional differences reflected between males and females. Compared to female controls (Fig. 4D showing 5 controls, 5 ME/CFS subjects), ME/CFS female CD4 and CD8 T cells show on average 26.5% and 28.6% higher proliferation respectively. Moreover, the regression lines in Fig. 4D show that CD4 and CD8 T cell proliferation among controls has 10x greater slope (blue line) with respect to ROS, compared to ME/CFS females (red). To check the generalizability of these findings, we evaluated in a separate experiment the relationship between ROS and T cell proliferation for 4 controls and 5 Long COVID female donors (Fig. 4E). Compared to controls, LC females on average show 39.7% and 42.8% higher CD4 and CD8 T cell proliferation, respectively. Consistent with the results in Fig. 4D, CD4 and CD8 T cell proliferation among controls has 10x greater slope (blue) with respect to ROS, compared to LC females. Together, these findings indicate that ME/CFS and LC T cells from female donors hyperproliferate, compared to controls, suggesting one potentially targetable source of patient fatigue.

In contrast, our analysis of 7 controls and 5 ME/CFS male donors found no differences in T cell proliferation on average (Fig. 4F). Comparing the slopes in the regression lines for Fig. 4F shows control male CD4 and CD8 T cell proliferation has 3.5 and 2.3x higher slope with respect to ROS, compared to ME/CFS males. While T cells from ME/CFS male donors do not hyperproliferate, their T cells exhibit greater insensitivity to ROS levels, compared to control counterparts.

These results suggest that the capacity for an individual ME/CFS T cell to proliferate is tuned differently or is insensitive to higher oxidative stress levels, pointing to a potential functional defect in ME/CFS T cell proliferation. As memory T cells account for the heterogeneity in the T cell glutathione profile and stimulation drives the formation of memory cells, our results imply deficient adaptive immune responses in ME/CFS donors. This finding agrees with an influenza vaccination study showing T cell hyperproliferation in the ME/CFS group versus controls, consistent with our result showing higher proportion of proliferating T cells upon in vitro stimulation^52^. Additionally, characterization of stimulated CD4 T cells from LC donors^12^ found significantly higher levels of intracellular IL-2 levels, which are primarily produced from T cells through ROS-NFAT signaling. While neither of these cited studies probed for sex-specific differences in T cell proliferation, our findings highlight that the sex-specific pathways in redox dysregulation also shape the functional differences in male and female ME/CFS or LC adaptive immune responses, where female T cells hyperproliferate and male T cells have an insensitive response to ROS.

As T cell proliferation is associated with a 10-fold increase in energy usage due to elevated protein synthesis^49, 50^, these findings also point to a potential source of patient fatigue especially in ME/CFS female donors, where T cell activation drives rampant proliferation in CFS donors. Based on this hyperproliferation and the specific characterized pathways in Fig. 2 and 3, we attempted to assess our findings in a clinical context and evaluated if ROS-modulating drugs could attenuate this hyperproliferation.

### 5. ME/CFS donor T cell hyperproliferation can be attenuated with metformin

We evaluated total ROS levels in three patients (27, 30, 31) from the family population-omics profiling (fPOP) cohort^53^, who presented with chronic symptoms (Figure 5A, table). The clinical ME/CFS diagnosis, based on NAM criteria^7^, was not available at the time of experimental characterization. Therefore, to contextualize changes in ROS levels, these donors were compared to 6 healthy controls (4 females, 2 males), along with samples from a ME/CFS presenting patient’s (patient 27) mother and father.

**Figure 5.**
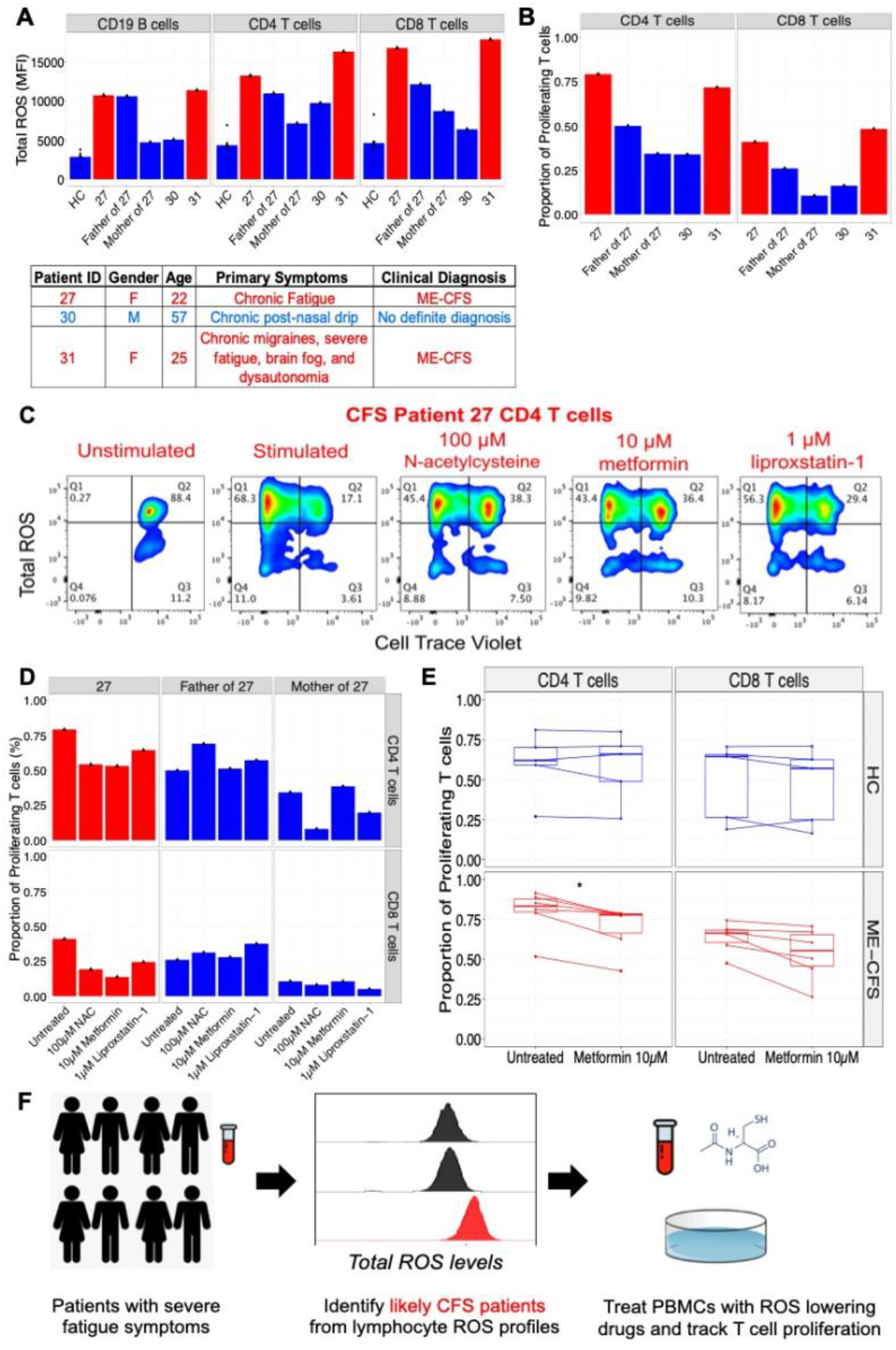
Identification of CFS patients using ROS signatures and tracking T cell proliferation enables personalized selection of ROS-lowering drugs to redress T cell hyperproliferation in CFS donors. (A) Study of fPOP patients flagged patients 27 and 31, along with the father of 27, based on measured total ROS levels (DCFDA) in lymphocytes. Patient clinical information shown in table identifies 27 and 31 as ME/CFS patients, based on the National Academy of Medicine ME/CFS diagnostic criteria^7^. (B) Using CellTraceViolet staining, the proportion of proliferating T cells 5-days after stimulation were assessed for fPOP donors. CFS donor CD4 and CD8 T cells with higher ROS are also associated with higher proportion of proliferation, consistent with Fig. 4E. Proportion of proliferating T cells were determined, based on same methods in Figure 4D. (C) To reduce proportion of proliferating T cells, PBMCs were treated at day 0 with ROS-modulating drugs, including N-acetylcysteine, metformin, and liproxstatin-1. Effects of drug on proliferation and ROS levels are shown 5-days after stimulation, where drugs reduce the proportion of high ROS proliferating T cells in Q1. (D) Comparison of drug effects between donor 27 and donor 27’s mother and father shows tested ROS-lowering drugs selectively modulate T cell proliferation in the CFS donor 27 and not in the parents. (E) Extension of metformin treatment to 5 additional HC and 5 ME/CFS donors finds a statistically significant reduction in the proliferation of ME/CFS CD4 T cells (p = 0.041). The drug does not alter proliferation in HC donors, highlighting its selectivity in ME/CFS donors. (F) Summary of Precision Medicine methodology for identifying ME/CFS patients and ROS-modulating drugs to address oxidative stress and hyperproliferation of T cells.

Comparison of total ROS levels in flow cytometry enabled us to flag patient 27, the father of 27, and patient 31, based on elevation of total ROS levels (Figure 5A). Overlaying our ROS data with patient symptoms showed that both patients 27 and 31 could be diagnosed as ME/CFS+, based on the NAM criteria^7^. Additionally, patient 30, who exhibited chronic post-nasal drip but did not report any symptoms of fatigue, was not flagged by our assay, highlighting its sensitivity in distinguishing between patients with chronic symptoms (Figure 5A). While our assay did not flag the mother of 27, our assay detected higher ROS levels in the father of 27 compared to healthy controls but still lower than patient 27.

As higher oxidative stress in T cells is associated with proliferation, we used CellTraceViolet staining to track the proliferation of fPOP donor T cells, upon anti-CD3/anti-CD28 and IL-2 stimulation. Expectedly, our findings showed a higher proportion of T cells from patients 27 and 31, who both presented with ME/CFS symptoms, exhibited elevated ROS levels and higher proliferation upon stimulation, compared to both parents of 27 and patient 30, who showed no fatigue symptoms and no elevation in ROS levels (Figure 5B). These differences were observed across both CD4 and CD8 T cells, where ∼1.8x and ∼2.4x higher proportion of patient 27’s CD4 and CD8 T cells respectively proliferated upon stimulation compared to her parents (CD4 T patient 27 - 79.3%, father of 27 – 50%, mother of 27-34.3%; CD8 T patient 27 – 41.0%, father of 27 – 26.0%, mother of 27 - 10.6%).

Based on the observation that ME/CFS donors have higher T cell ROS levels, associated with significantly higher proportion of proliferating T cells, we assessed whether ROS modulating drugs could lower the proportion of proliferating T cells. From our metabolic characterization (Figure 2, 3), we tested three drugs, including N-acetylcysteine, metformin, and liproxstatin-1. N-acetylcysteine (NAC), an FDA approved drug, helps replenish glutathione levels, thereby acting as a critical reducing agent/anti-oxidant to combat oxidative stress. Metformin, also an FDA approved drug, can inhibit mitochondrial complex I proteins and subsequently induce SOD2 expression, thereby reducing ROS formation^54^. Our earlier assays identified SOD2 loss as a characteristic of LC and some ME/CFS donor lymphocytes (Figure 2C) and double-blinded phase III studies on >1300 patients have shown metformin reduces Long COVID incidence by 41%, especially among female subjects. Based on these results, metformin was also tested for its capacity to curtail T cell hyperproliferation in ME/CFS donors. Finally, as our lipid metabolic studies showed GPX4 changes and higher lipid peroxidation in ME/CFS and LC donors, we tested liproxstatin-1^56,57^, a potent GPX4 modulator that can slow the accumulation of lipid peroxides.

Among the fPOP donors, we treated patient 27, 27’s father, and 27’s mother PBMCs at day 0 and compared the T cell proliferation after 5 days of stimulation between conditions. Comparing the stimulated conditions with treatment to the untreated for patient 27, our flow cytometry data suggested that all three drugs could modulate the proportion of high ROS proliferating T cells (Figure 5C, Q1). Specifically, compared to 79.3% of proliferating CD4 T cells upon stimulation, NAC, metformin, and liproxstatin-1 treatment were able to lower this proportion to 54.3%, 53.2%, and 64.5% respectively. Moreover, we found that patient 27’s T cell proliferation was uniquely responsive to these ROS-lowering drugs (Figure 5D). In contrast, both patient 27’s mother and father CD4 and CD8 proportion of proliferating T cells did not significantly change in response to any of these drugs (Figure 5D).

To test the generality of these findings, we tested these drugs on 5 HC and 6 female ME/CFS donors, including those reported in Figures 4D. While our assays found that NAC did not have any effect in reducing T cell hyperproliferation in ME/CFS or LC female donors (Supplementary Figure S6, ME/CFS CD4 p = 0.82, CD8 p = 0.7; HC CD4 p = 1, CD8 p = 0.42; Supplementary Figure S7, LC CD4 p = 0.69, CD8 p = 0.55), metformin had a statistically significant effect in reducing CD4 T cell hyperproliferation in ME/CFS female donors (Figure 5E, CD4 9.8% reduction in proliferation p = 0.041, CD8 10.5% reduction in proliferation p = 0.39). As NAC increases GSH levels, which are already significantly elevated in ME/CFS donors, our assays suggest that increasing glutathione levels is not sufficient to address the oxidative stress in ME/CFS or LC donor lymphocytes. Also, our analysis on these donors showed that metformin uniquely reduced the proliferation in ME/CFS donor T cells with no effect among the healthy controls (Figure 5E, HC CD4 p = 1, CD8 p = 0.69). While liproxstatin-1 did not show a statistically significant effect in lowering T cell proliferation among female CFS or LC donors (Supplementary Figure S6, S7), the results again show that ME/CFS donor T cell proliferation is uniquely modulated by the drug and not healthy controls (Supplementary Figure S6, ME/CFS CD4 p = 0.31, CD8 p = 0.18; HC CD4 p = 1, CD8 p = 0.69). Metformin treatment had a smaller and non-statistically significant effect in LC female PBMCs (Supplementary Figure S7B, CD4 p = 0.31, CD8 p = 0.42), where treatment reduced on average CD4 and CD8 T cell proliferation in female LC donors by 5.2% and 7.6% respectively.

Consistent with the finding that male ME/CFS T cells did not hyperproliferate and displayed greater insensitivity to ROS, we did not observe any reduction in T cell proliferation upon treatment with NAC, metformin, or liproxstatin-1 (Supplementary Figure S8). Regardless, these findings demonstrate the *possibility* of a Precision Medicine approach for helping potentially diagnose and treat ME/CFS (Figure 5F), especially among females. As shown in our study, tracking redox aberrations in lymphocytes can be used to identify individuals, who may present as ME/CFS based on clinical symptoms. Moreover, the selectivity of metformin and liproxstatin-1 in ME/CFS donors underscores that modulating oxidative stress is a tunable molecular link for controlling the extent of T cell proliferation in ME/CFS donors. It follows that tracking T cell proliferation in the context of ROS levels can be pinpoint which individuals may potentially benefit from ROS-lowering drugs and even identify novel drug candidates that mediate this link between ROS and T cell proliferation.

## Discussion

While ME/CFS and Long Covid share many clinical features along with other post-acute infection syndromes^6^ and are increasingly diagnosed together, there are no approved molecular diagnostics or treatments for these patients. But previous studies have hypothesized oxidative stress as a common biochemical signature^26^ and even shown some evidence in circulating serum redox proteins (e.g., myeloperoxidase)^59, 75^ among LC patients. Unique to our work, the analysis of intracellular pathways shows that both diseases share signatures of elevated oxidative stress, specifically among lymphocytes, compared to healthy controls that would impact mitochondrial function. Additional intracellular metabolic characterization using a variety of approaches shows that these changes are reflected in multiple oxidative stress pathways, including alterations in glutathione, superoxide dismutase, lipid oxidative damage, etc. Also, our work is the first (as far as we know) to pinpoint a potential targetable cellular mechanism, where these metabolic changes clearly affect lymphocyte activation and likely the immune response generally, and if systemic could explain the prevalence of fatigue and other symptoms in these two syndromes.

Our findings also show sex-specific redox signatures in ME/CFS patients, such as elevated total ROS levels in women versus men, suggesting that the pathophysiology for ME/CFS and LC are distinct between genders. Published studies in LC and ME/CFS have shown redox abnormalities in circulating components^75^, including serum proteins^59^ and erthryocytes^60^. While additional studies are needed to understand the mechanistic basis of these sex-specific differences in ME/CFS and LC, we offer one potential explanation. As sex hormones such as estradiol can regulate the serum levels of anti-oxidant enzymes, including catalase, glutathione peroxidase, and SOD2, the distinct underlying redox biology for ME/CFS may be partially explained by differences in these hormone levels^58^. Moreover, estrogen also shapes the activation of CD4 T cells, which express the estrogen receptor (ERα)^62^. Comparison of in-vitro stimulation of T cells between males and females shows higher expression of anti-viral genes in females^64^ (IFNG, RIGI, OAS1) and higher production of IFNγ upon stimulation^62^. Additionally, both our study and a recent deep phenotyping study^65^ on ME/CFS patients show differences in fatty acid oxidation pathways between ME/CFS males and females, with the results here uniquely highlighting larger changes in lipid peroxidation. As androgens modulate peroxisome proliferator-activated receptor (PPAR)α levels in CD4 T cells, where PPARα critically regulates the production of fatty acid oxidation enzymes, sex differences in PPARα CD4 T cells may explain this difference in lipid metabolism^66,67^. Next, our findings suggest that these metabolic differences likely impair the adaptive immune response in ME/CFS and LC individuals. Through studying the extreme value distributions in the glutathione profiles, we find that memory CD4 T cells are critical contributors to the heterogeneity in bulk CD4 population profiles. These findings, show that CD4 T cells oxidative profiles in LC patients exhibit a long-tail phenomena, also caution against solely comparing medians/means between donors, which does not capture this heterogeneity. By comparing T cell proliferation upon antigen stimulation, we find oxidative stress linearly scales with the proportion of proliferating T cells, meaning a higher fraction of female CFS donor T cells proliferate upon stimulation. In addition, our results showing altered sensitivity of T cell proliferation to oxidative stress levels among CFS donors (compared to HCs) carries additional implications. With the higher energy demands needed for T cell proliferation, these differences suggest the availability of energy as a significant and unique contributor to fatigue in CFS donors. We suggest that this lymphocyte dysfunction, driven by oxidative and mitochondrial damage, acts as an “energy sink”, much as an active infection does, draining the body of available energy and giving rise to debilitating fatigue and other sequelae. Moreover, it suggests that at least in females, ROS levels may serve as a tunable link for adjusting T cell proliferation, with CFS donor T cells showing insensitivity to higher ROS levels.

Third, as demonstrated in the fPOP study^53^, our work demonstrates a potential Precision Medicine methodology to identify specific pre-existing FDA approved and novel candidates that can adjust ROS levels and subsequently curtail T cell hyperproliferation in CFS donors. In light of a clinical study that has shown metformin’s efficacy in lowering symptoms of Long COVID in 41% of treated subjects^55^, especially in female subjects and those with higher BMI, our results offer one plausible mechanism of action in LC patients and more broadly suggest the link between oxidative stress and aberrant T cell proliferation can be exploited to identify novel drug candidates.

Although our findings were focused on identifying a conserved signature between ME/CFS and LC patients, based on commonly presented symptoms, our analysis suggests several important differences. For example, while both ME/CFS and LC donors show elevated ROS levels in T cells, the specific aberrant anti-oxidant pathways appear to differ slightly, where LC patients overall show lower mitochondrial superoxide dismutase and calcium levels compared to ME/CFS donors. Additionally, our comparison of the extreme value distributions found that LC T cells show a heavy-tailed oxidative stress profile and CFS donor T cells show a shifted but bounded extreme value distribution. As one difference between these groups in our study corresponds to the duration of symptoms, it is possible that LC donors capture an *intermediate state* between HC and ME/CFS donors. One interpretation of this concept is that continuous exposure to high oxidative stress in lymphocytes may cultivate tolerance to ROS, as a possible protective mechanism for some patients^63^. This tolerance may consequently lead to insensitivity to ROS levels, upon stimulation, as evidenced by our findings in ME/CFS T cells.

These results also raise the question as to how these oxidative stress pathways are triggered and maintained. Here we can only speculate that since Long Covid is clearly triggered by an infection, the initial elevation of ROS levels necessary for lymphocyte activation is overly prolonged in some individuals, perhaps caused by an impaired ability to clear the virus, causing damage to mitochondrial membrane physiology. This ROS elevation may then return to baseline (or get cleared more rapidly) in males but not females, but in both sexes with the LC syndrome the damage is long lasting. In ME/CFS it has long been thought that there was a causative infection, although a specific pathogen has not been identified, despite intensive efforts. Nevertheless, it has been noted that other infections have produced LC-like symptoms in particular individuals^6^, so the pathologies we document here might be a feature of other infections as well. It may be that individuals with sub clinical immune deficiencies suffer from a more prolonged infection^70^ and accompanying ROS elevation that produces lasting mitochondrial damage.

Additional studies in larger and diverse cohorts will be needed to test the generality of these conclusions further. Regardless, we are hopeful that the identified pathways from our findings may provide a blueprint to guide the development of possible diagnostics and therapies to help ME/CFS and LC patients.

## Acknowledgments

VS acknowledges discussions with Davis & Mischel lab members, along with the help of Huijun Yang and Rick Cuevas. We thank the shared FACS facility for assistance in flow cytometric analysis and cell sorting (Symphony 1S10OD026831-01, LSRII.UV S10RR027431-01, PICI Purchased by Parker Institute for Cancer Immunotherapy).

## Competing Interests

VS, SS, MMD, PM, HB, and MS are inventors on a patent related to oxidative stress signatures in ME/CFS and LC. SS co-founded and is a scientist at Material Alchemy (MA), an independent entity for Designing Materials for Sustainability. Turium was developed by MA for analyzing complex systems chemistry and is available to academia for research with licensing.

## Funding

We are grateful for funding by the National Institute of Allergy and Infectious Diseases (NIAID) (U19-AI057229, 5R01AI139550) and the Howard Hughes Medical Institute to MMD. We also acknowledge funding from the Khosla family gift fund.

