## Supplemental Text & Figures for "Oxidative Stress is a shared characteristic of ME/CFS and Long COVID"

### Methods Description
### Patient Description:

For the current study, we selected three cohorts (Figure 1A, main text).

- Healthy control (HC) samples were obtained from the Stanford Blood Bank under IRB-40146. HC participants must be 17 years of age or older and complete a medical history questionnaire that includes recent symptoms related to COVID-19 or a positive test two weeks before the blood donation and test negative for hepatitis B and C, HIV, HTLV, syphilis, and West Nile Virus (https://stanfordbloodcenter.org/medical-history-questionnaire/).
- ME/CFS patients were eligible for inclusion in the study under IRB-40146 if they had been diagnosed using the National Academy of Medicine and Canadian Consensus Criteria (CCC). Severity was assessed using Bell's disability scale. Informed consent was obtained through REDCap, and study data were collected and managed using REDCap electronic data capture tools hosted at the ME/CFS Collaborative Research Center at Stanford University.
- Long COVID-19 participants were patients from the Stanford Post-acute COVID syndrome (PACS) clinic. These patients met the CDC case definition of Long COVID[17], must be 18 years of age or older, must have new symptoms after SARS-CoV-2 infection and evidence of positive infection (positive PCR, antigen test, or antibodies before SARS-CoV-2 vaccination). Blood collection, demographic, and clinical data were collected from this study cohort under IRB approval protocol (IRB-64344) and this information was blinded to the bench researchers. Based on the National Academy of Medicine criteria, 9/15 (60%) met the diagnostic criteria for ME/CFS.

| | Clinical Cohorts Demographics | | |
|---|---|---|---|
| Clinical Cohorts | Healthy Controls (HC) | Chronic Fatigue Syndrome (ME/CFS) | Long COVID (LC) |
| Number of Participants | 16 | 15 | 15 |
| Mean age | 53 | 52 | 47 |
| % female | 46.6% | 76.9% | 53.3 |
| % ME/CFS | N/A | 15/15 (100%) ** | 9/15 (60%) * |

*IOM criteria, **IOM and CC criteria, N/A Non applicable

### PBMC Preparation:

Human primary peripheral blood mononuclear cells (PBMCs) were isolated from the whole blood of both healthy donors and ME-CFS patients using SepMate™ 50 mL Tubes, following the protocol specifications. After isolation, cells were frozen in FBS + 10% DMSO and stored in liquid nitrogen tanks.

PBMCs were thawed and washed in pre-warmed RPMI media with benzonase. RPMI media included RPMI-1640 media supplemented with 10% dialyzed FBS, 1% glutamax, and 1% penicillin/streptomycin. After spinning down cells at 400g for 5 minutes, cells were counted and viability was evaluated by Trypan blue staining. Cells were passed through a 70µm cell strainer, spun down at 400g for 5 minutes, and re-suspended at 1-2 million cells/well in RPMI media. After resting PBMCs overnight for 12-18hrs at 37C, cells were stained the following morning with metabolic dyes.

**Staining:** To stain for mitochondrial parameters, cells were stained with several metabolic dyes.

Cells were then collected, spun down 2x at 400g for 5 minutes, stained with live-dead dye and Fc receptor (FcR) block in FACS buffer (PBS, 10% dialyzed FBS, 1mM EDTA). After 10 minutes, cells were subsequently stained with surface antibodies for 30 minutes. Due to the variety of metabolic dyes used, we also set-up multiple panels accordingly. We list all antibodies used across any panel. Antibodies included anti-CD3 (BUV805-UCHT-1, BV605-OKT3), anti-CD8 (BUV395, RPA-T8;

BUV737, RPA-T8), anti-CD4 (BV650, RPA-T4, AlexaFluor700, RPA-T4), anti-CD19 (PerCP/Cy5.5, HIB19). For single colored controls, combination of compensation beads and stained cells were used for surface antibodies and metabolic dyes, respectively. Naïve and memory CD4 T cells were stained using anti-CCR7 (PerCP/Cy5-5, Clone G043H7; BV421, Clone G043H7) and anti-CD45RO (PE/Cy7, UCHL1, BioLegend) antibodies. Activated T cells were stained in proliferation assays using anti-CD69 (BUV395; Clone FN50) and anti-CD137 (BV750, Clone-4B4-1). Flow cytometric and fluorescence activated cell sorting analysis was performed on an BD Aria Fusion sorter and BD FACS Symphony A5 instruments. Flow cytometry data was analyzed using FlowJo software (BD). Staining levels were compared using median, mean , and 95$^{th}$ percentile fluorescence intensity (MFI = median fluorescence intensity).

**Intracellular Staining for SOD2:** After staining PBMCs with surface markers and live-dead dyes, cells were fixed and permeabilized, using BD Cytofix/Cytoperm kit. Subsequently, cells were stained with antibodies to intracellular proteins (SOD2 (FITC, Clone: 3A6C2, ThermoFisher) for 45 minutes at 4C.

**Immunofluorescence Staining Microscopy:** Upon CD3 (Alexa594, UCHT1, BioLegend) and GPX4 staining (ThermoFisher, Catalog ID: PA5-102521), stained cells were transferred to microscope slides, using cytospin preparation. Images were acquired using a Leica DMi8 Thunder Imager equipped with a Leica DFC9000 GT Camera.  Z-stack images were rendered and segmented in three-dimensional space using AIVIA Pixel Classifier and Recipe Analysis for subcellular signal quantification.

**Proliferation Assays:** PBMCs were labeled with proliferation dye according to manufacturer's instructions using CellTrace Violet (ThermoFisher). After resting labeled PBMCs for an hour, cells were stimulated with anti-CD3/anti-CD28 antibodies (StemCell Technologies) with IL-2 50 IU/mL. The extent and proportion of proliferating T cells were compared 5 days after stimulation.

**Mass Spectrometry:** First, 200,000 CD3 T cells were sorted, pooled across several healthy control and ME-CFS donors. Metabolites were immediately extracted from cells using 80% methanol and 20% water mixture with mass spectrometry grade internal standards (Ref. 23, main text). The cells are repeatedly vortexed and sonicated, upon which the proteins are precipitated, and extracted metabolites are dried under nitrogen stream and stored in -80C until ready to run. The hydrophilic interaction (HILIC)-MS analysis was conducted using a Thermo UltiMate 3000 UHPLC coupled to a Thermo Q Exactive HF mass spectrometer. HILIC experiments were performed using a ZIC-HILIC column (2.1 × 100 mm, 3.5 µm, 200 Å; Merck Millipore) with mobile phase solvents consisting of 10 mM ammonium acetate in 50/50 acetonitrile/water (A) and 10 mM ammonium acetate in 95/5 acetonitrile/water (B). Peaks were assigned by matching m/z and retention time with external databases and pre-purchased analytical standards. The fragmentation profile for all peaks were compared with several databases, including HMDB, METLIN, Lipid MAPS, and MassBank to assign the peak identities.

**Turium:** This is a system chemistry tool developed for complex chemical analysis. After the identification of analytes, mass spectrometry data was processed for Turium analysis, where metabolic networks were constructed of healthy controls and ME-CFS patients. From 45,507 unique detected analytes, 747 were identified based on comparisons with known analytical standards (Ref. 23, main text), and 327 different metabolic pathways were mapped. The resulting specific metabolic reactions were checked with KEGG Pathway Database (https://www.genome.jp/kegg/). The *difference* between the metabolic pathways, those in ME-CFS donors but not in healthy controls, was computed and visualized using networkx package in R v3.6.3.

**Supplementary Figure S1**

(A) Sample gating strategy is shown for CD19 B cells, CD4 and CD8 T cells.

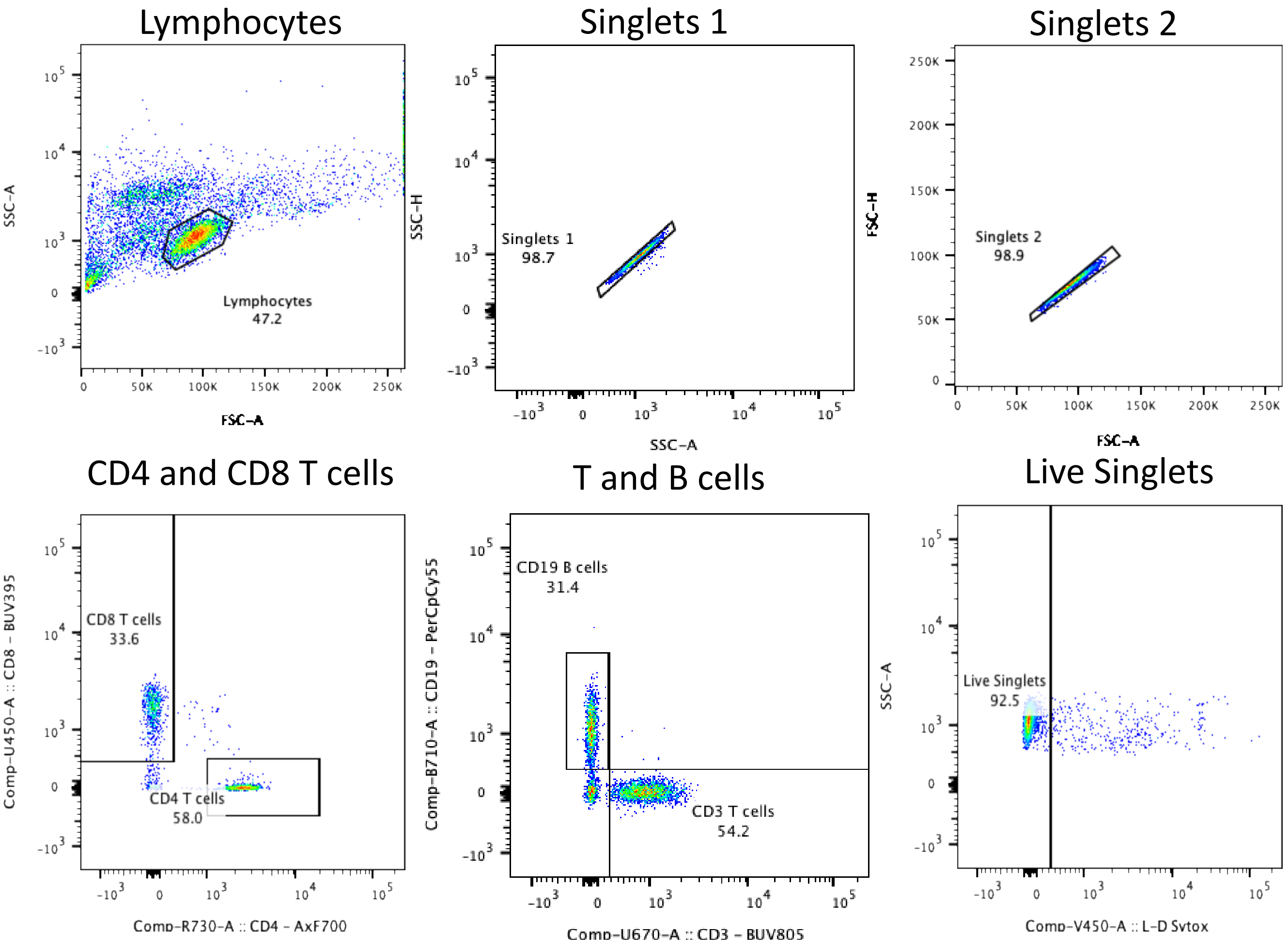


(B) After gating for CD4 T cells, CCR7 and CD45RO surface markers are used to define memory and naïve populations.

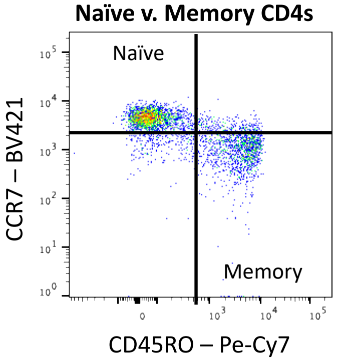

**Supplementary Figure S2.** Comparison of mitochondrial parameters between healthy controls (HC), ME-CFS, and LC donors. While ME-CFS and LC donors appear to present with lower mitochondrial ATP across lymphocyte populations compared to HC donors, no differences in mitochondrial mass or membrane potential are observed.

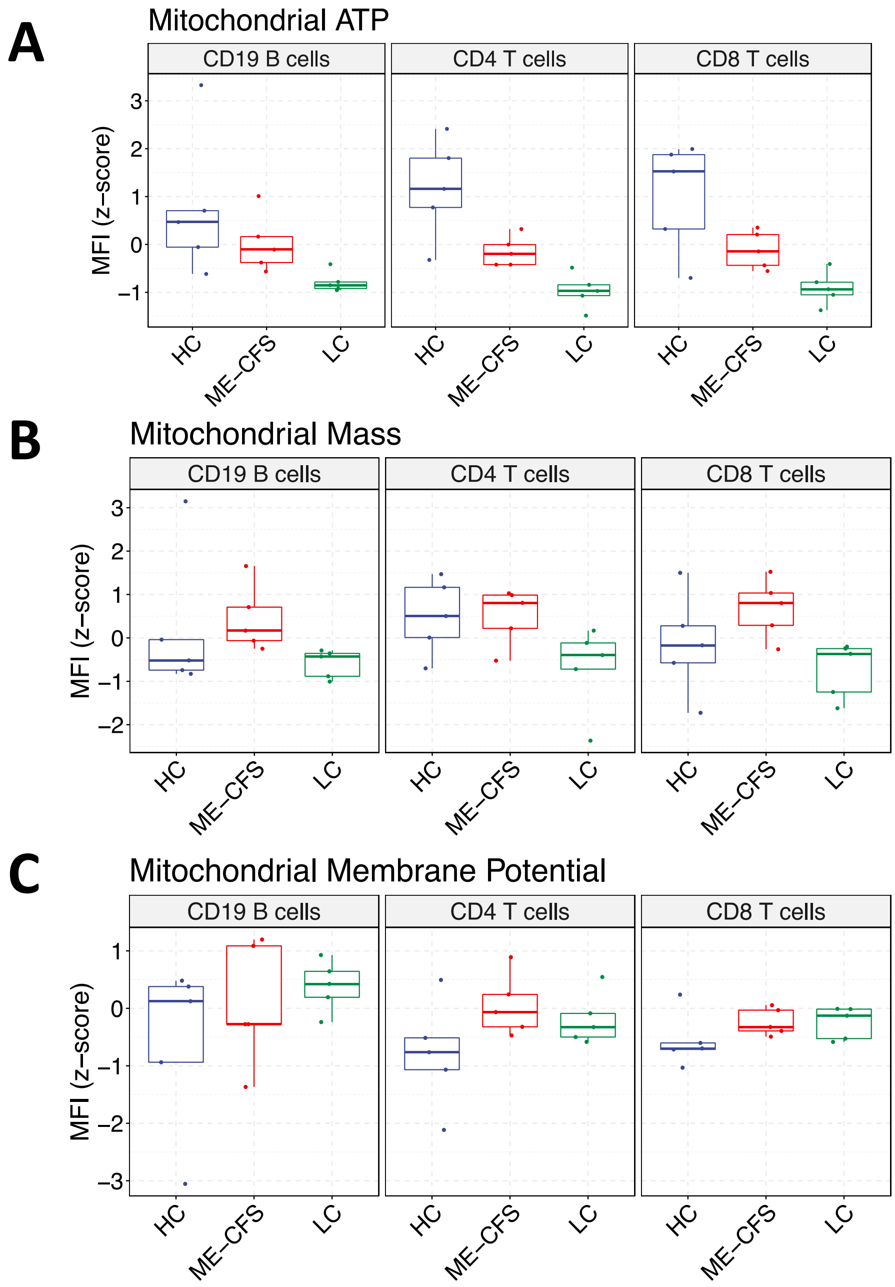

**Supplementary Figure S3**. Comparison of total ROS levels based on patient characteristics

(A) No difference in Total ROS levels (MFI) by fatigue severity

(B) Association between Total ROS levels (MFI) and duration of LC symptoms

(C) No association with age

(D) Association with BMI in Long COVID group, where donors with higher BMI have higher total ROS levels

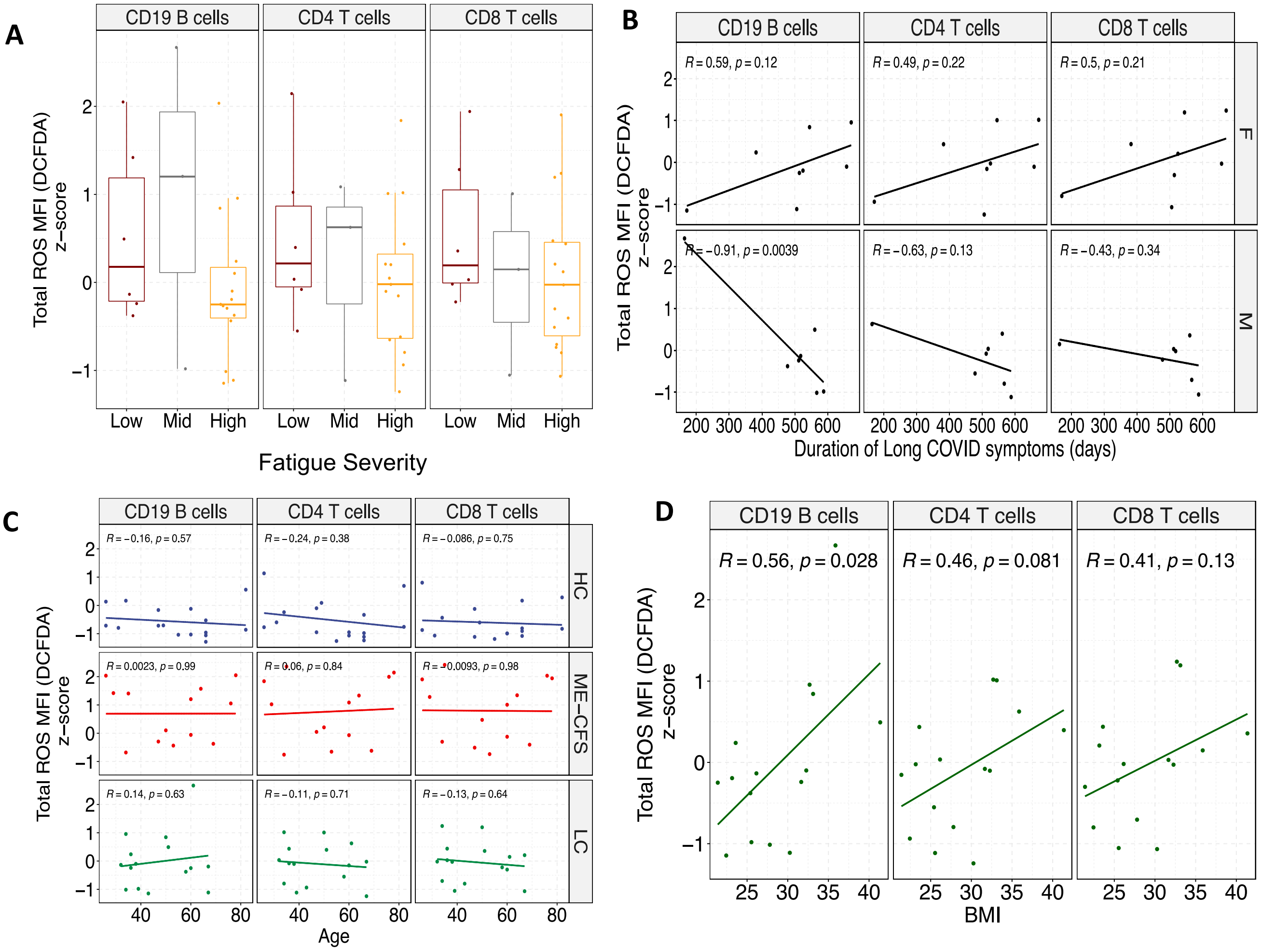

**Supplementary Figure S4.** Comparison of redox parameters between HC, ME-CFS, and LC donors across males and females

(A) Comparison of mitochondrial calcium levels identifies statistically significant differences uniquely in females and not in males (two-sided t-test: females: CD4T ME-CFS p = 0.0063, CD4T LC p = 0.0172, CD8 T CFS p = 0.0137, CD8T LC p = 0.24, CD19 B CFS p = 0.0024, CD19 B LC p = 0.0097).

(B) LC donors across both males and females show decreases in SOD2 levels, compared to HCs of the same gender

(C) Comparison of lipid peroxide levels are shown, where lower y-axis values correspond to higher levels of lipid peroxides. ME-CFS and LC donors across genders exhibit similar levels of lipid peroxides. As HC males have lower levels of lipid peroxides compared to HC females, the differences in lipid peroxidation are more pronounced.

(D) Comparison of lipid droplet levels are shown between groups, where ME-CFS and LC donors show decreases in lipid droplets compared to HCs across both genders. Similar to lipid peroxide comparisons, HC males have higher levels of lipid droplets compared to HC females, meaning differences in ME-CFS and LC groups are greater for males.

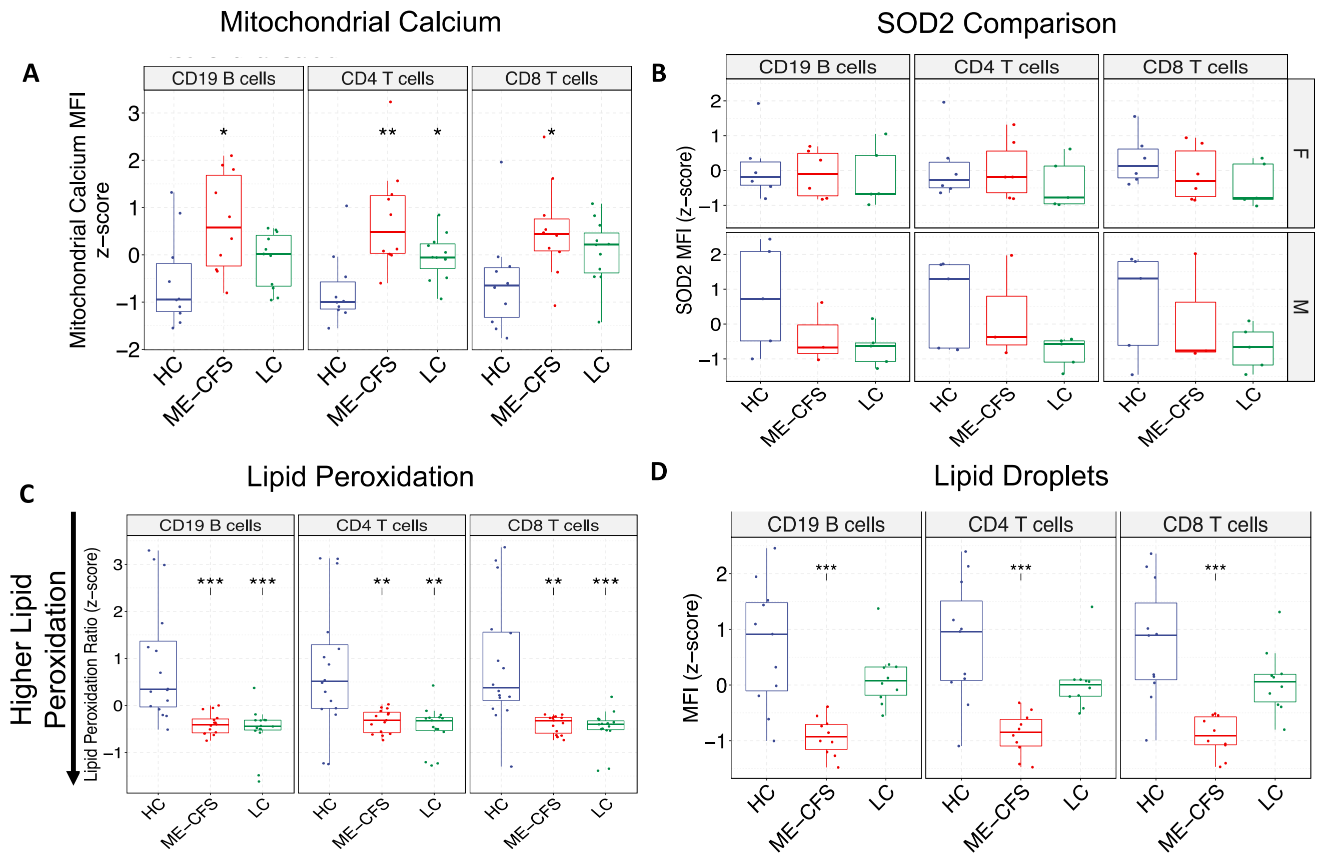

**Supplementary Figure S5.** Comparison of CD69, CD137 MFI levels between HC and ME/CFS CD4 and CD8 T cells. These differences are shown 5-days post-activation with anti-CD3/anti-CD28 antibodies and IL-2.

(A, B) Plots are shown for female HC and ME/CFS donors.

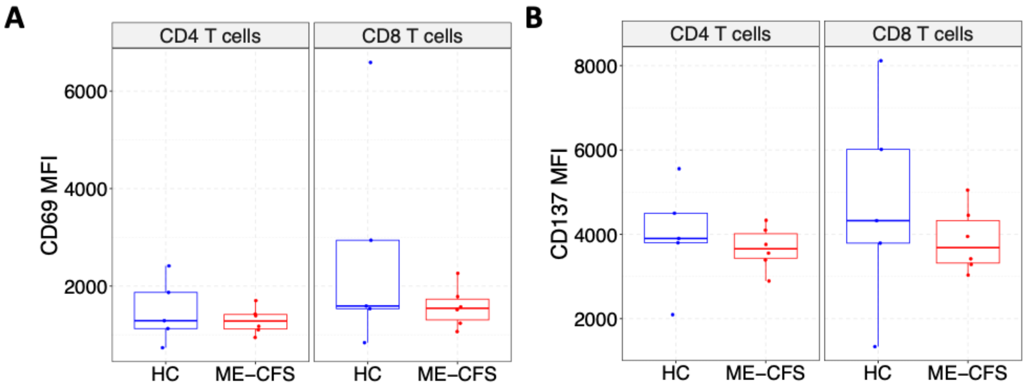


(C, D) Plots are shown for male HC and ME/CFS donors

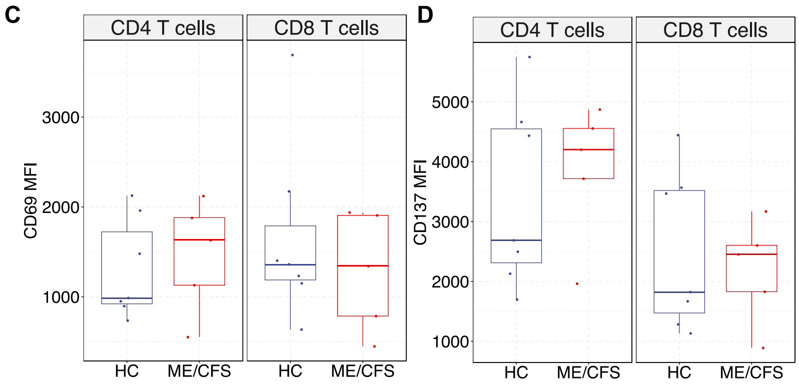


(E, F) Plots are shown for female HC and LC donors

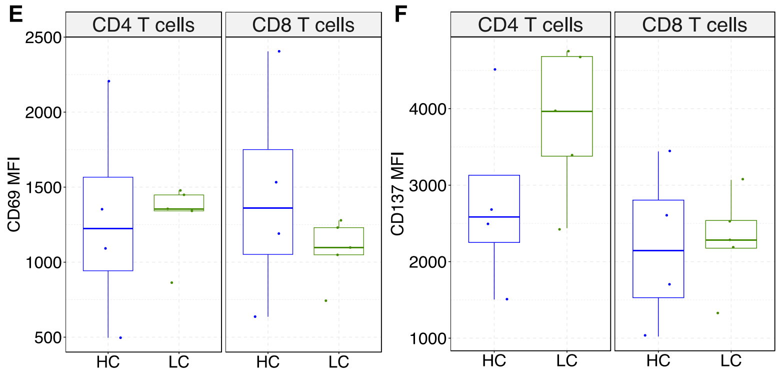

**Supplementary Figure S6.** Treatment of HC and female ME-CFS PBMCs with (A) 100 µM N-acetylcysteine and (B) 1 µM liproxstatin-1
The proportion of proliferating T cells is shown 5-days post activation for 5 HC, 6 ME/CFS donors. While no difference or decrease in proliferating T cells are observed upon NAC treatment (A), liproxstatin-1 treatment, which targets GPX4, selectively lowers proliferation in ME-CFS and not the HC donors.

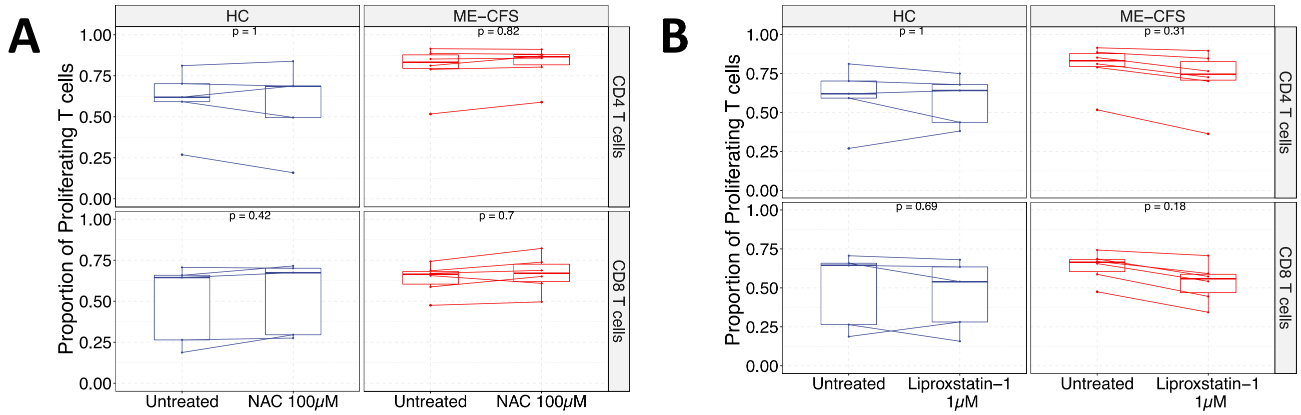

**Supplementary Figure S7.** Treatment of HC and female LC PBMCs with (A) 100 µM N-acetylcysteine (B) 10 µM metformin (C)1 µM liproxstatin-1

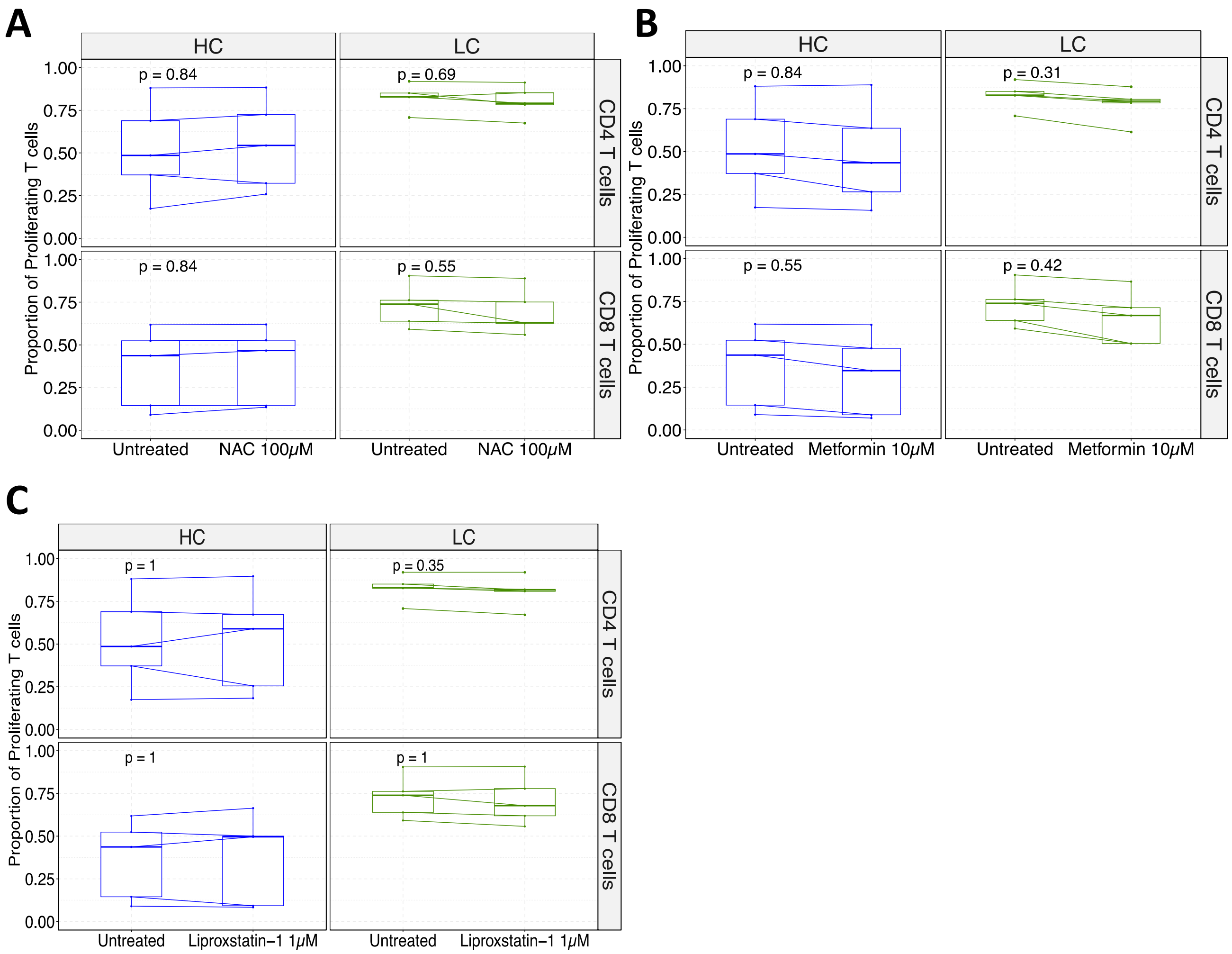

**Supplementary Figure S8.** Treatment of HC and male ME/CFS PBMCs with (A) 100 µM N-acetylcysteine (B) 10 µM metformin (C)1 µM liproxstatin-1

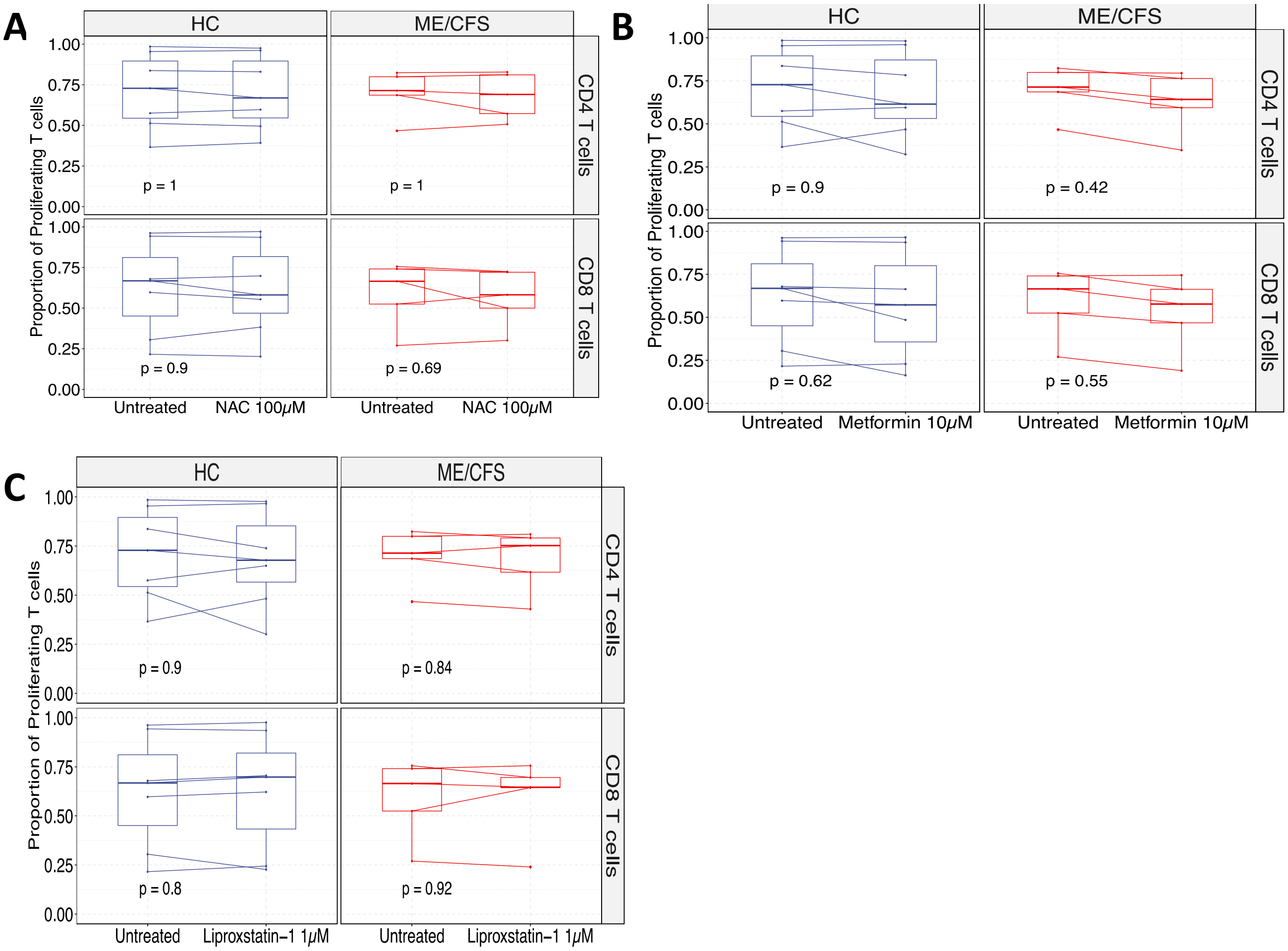
